# Single-locus chromatin memory enables flexible spatial fate specification in the *Drosophila* visual system

**DOI:** 10.64898/2026.08.09.743789

**Authors:** Yen-Chung Chen, Sophia Lera Nielsen, Ben Jiwon Choi, Asif Bakshi, Rose Coyne, Mehmet Neset Özel, Claude Desplan

## Abstract

Spatial patterning generates neuronal diversity by compartmentalizing progenitors into domains with distinct molecular identities. However, these patterning cues are often transient in neurogenic domains, raising the question of how spatial information can be preserved and how rigidly it constrains neuronal fates. Here we show that a single-locus chromatin memory in the medulla of the *Drosophila* optic lobe faithfully preserves spatial identity while conferring the flexibility for specific neuronal fates to bypass spatial restriction. Medulla progenitors are partitioned into three spatial domains marked by Vsx1, Optix and Bifid. Domain-resolved single-cell multiome profiling reveals that progenies from different neuroepithelial domains are nearly indistinguishable for both transcriptome and chromatin accessibility, although persistent domain-specific accessibility in Vsx and Bi progenies is restricted to a single corresponding spatial-factor locus, *Vsx1*/2 or *Bifid*, respectively. These same factors are absent when neuroepithelial cells are converted to neural stem cells, but they are re-expressed in postmitotic neurons to execute domain-specific fates. Because this bookmarking is so restricted, specific classes of neurons can bypass spatial restriction by overriding only this single-locus chromatin state, adopting equivalent fates regardless of spatial origin. Other neurons ignore domain boundaries by expressing *Vsx1/2* through a program independent of their domain of origin. PRC2-mediated silencing restricts *Vsx1* re-expression to its home domain, while temporal identity and Notch signaling in newborn neurons determine which neurons engage or bypass the spatial program. Thus, single-locus chromatin memory preserves spatial information without making it an obligatory determinant of every neuronal fate.

## Introduction

Spatial patterning is a conserved strategy for generating neuronal diversity. In both invertebrate and vertebrate nervous systems, morphogen gradients and transcriptional cross-repression subdivide neural progenitors into molecularly distinct domains, each of which gives rise to specific neuronal types. The spatial transcription factors that define these domains are frequently transient, but spatial information is typically relayed to downstream effectors before the original factors are turned off. In the *Drosophila* ventral nerve cord, orthogonal columnar and row genes specify the spatial identity of neural stem cell (neuroblast) in a combinatorial fashion in each segment. Although their expression fades during neurogenesis^1–8^, relay factors or genome-wide chromatin priming bridges this transient spatial specification to the terminal selectors that define neuronal identity^9–13^. In the vertebrate neural tube, morphogen gradients induce transcription factors (TFs) that define progenitor domains. These TFs activate terminal selectors before expression of progenitor TFs diminishes^14^. In each case, a gene regulatory handoff ensures that spatial information is continuously transmitted from progenitors to neurons, even in the absence of the original patterning factors in neurons.

While spatial patterning enables the generation of diverse neuronal types, it also imposes a numerical tradeoff: subdividing a progenitor pool into smaller domains limits the output of each domain-restricted fate^15^. A neuronal type serving a repeated structure must be produced in the high numbers that its structure demands, and production across multiple spatial domains is one of several routes to meeting that demand. As a consequence, many neuronal types ignore spatial information rather than being restricted to a single domain, indicating that this constraint can be selectively bypassed.

The *Drosophila* medulla, a visual processing center comprising approximately one hundred neuronal types in different stoichiometries, provides a system in which both the mechanisms and the limits of spatial patterning can be examined. The medulla is organized into ∼800 repeated retinotopic columns, and multiple neuronal types occur at one neuron per column, which reflects a need for dense spatial sampling across the visual field. Within the outer proliferation center, neuroepithelial cells that generate medulla neurons are divided into spatial domains marked by distinct transcription factors along the antero-posterior axis, including anterior *Vsx1*, middle *Optix* and posterior *Bifid* (*Bi*)^16–19^ (**Figure 1A**). Following the conversion of neuroepithelial cells (NE) into neuroblasts (NB), a shared temporal TF cascade further diversifies progenitor identity within each domain^20–24^. Finally, Notch-mediated asymmetric division of an intermediate progenitor, or ganglion mother cell (GMC), generates two sibling neurons with distinct fates^20,21^ (**Figure 1B**). Together, the intersection of spatial, temporal, and Notch-dependent axes generates the diversity of medulla neuron types (**Figure 1C-D**)^25,26^. This combinatorial logic, however, is not universal: some neuronal types acquire the same identity regardless of the spatial domain in which they are born^16,27^.

**Figure 1.**
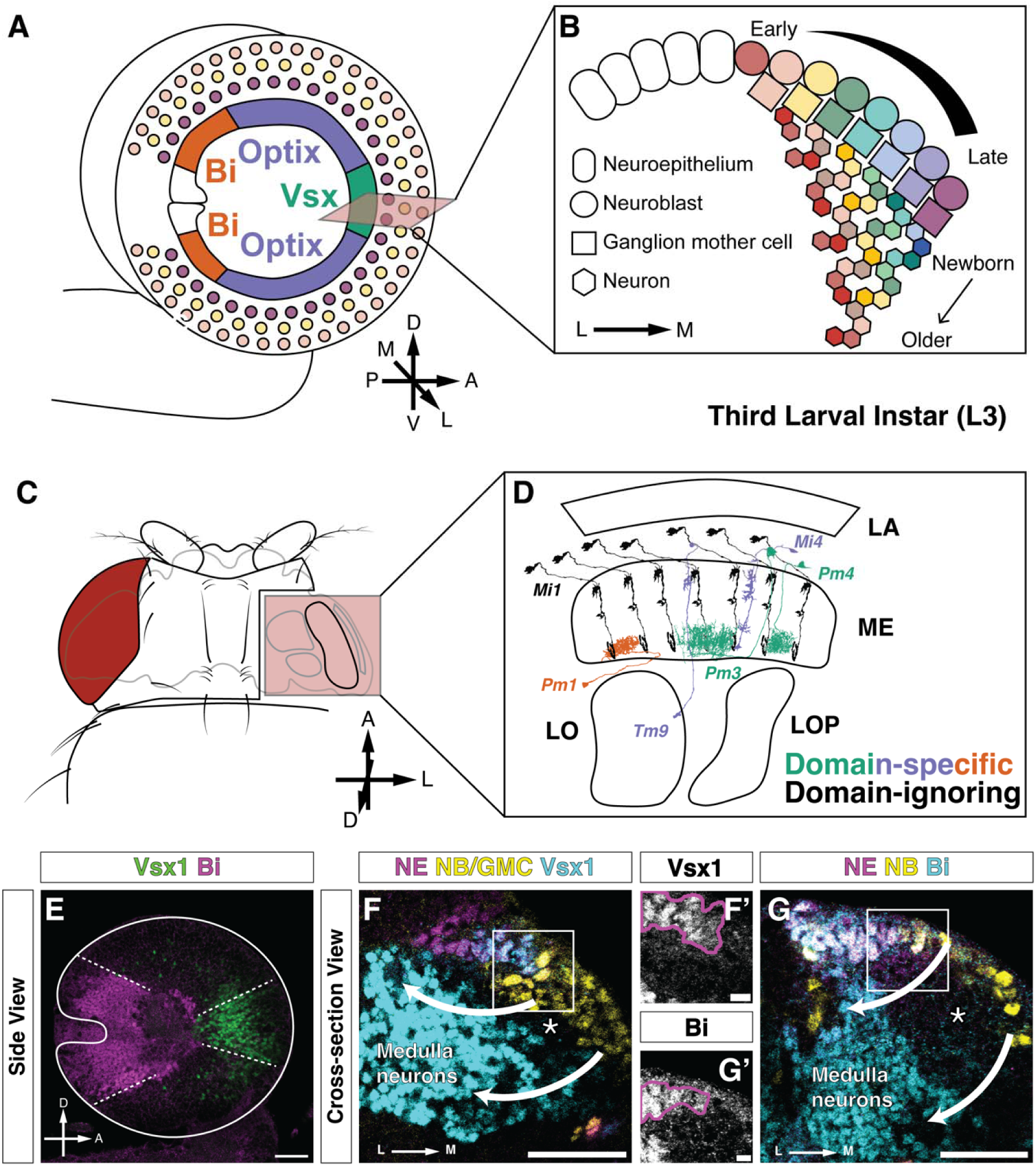
Spatial factors are transiently lost and later re-expressed during medulla neurogenesis. A, Side-view schematic of the third-instar larval optic lobe. Neuroepithelial cells of the outer proliferation center (OPC) are subdivided into spatial domains, while their neuronal progenies are further diversified by temporal identity. B, Cross-sectional schematic of OPC neurogenesis. Neuroepithelial cells convert into neuroblasts, which sequentially progress through temporal transcription factor windows and generate neurons through ganglion mother cell intermediates. Curved triangle: Early-to-late progression of neuroblast temporal cascade. Arrow: the direction of lineage extension pointing toward older/early-born neurons. A: anterior; P: posterior; V: ventral; D: dorsal; M: medial; L: lateral. C, Schematic of the adult fly head with the right optic lobe shown in the inset in D. D, Horizontal view of the adult optic lobe showing representative medulla neuron morphologies colored by spatial origin (green: Vsx; purple: Optix; orange: Bi; black: domain-ignoring Mi1 neurons). Colored neurons from right to left by soma position: Pm1 (Bi), Tm9 (Optix), Pm3 (Vsx), Mi4 (Optix), and Pm4 (Vsx). E, Side view of the larval optic lobe showing domain-enriched expression of Vsx1 (green) and Bi (magenta) in neurons of the medulla cortex. Dashed lines: approximate domain boundaries. The white lines mark the outline of the optic lobe. A: Anterior; D: Dorsal. Scale bar: 30μm. F, Cross-sectional view (as in B) of Vsx1 expression relative to Grh-expressing neuroepithelial cells (magenta) and Ase-expressing neuroblasts and ganglion mother cells (yellow). Vsx1 (cyan) is detected in the neuroepithelium and differentiated neurons, but not in the intervening progenitor zone. Asterisk: nascent neurons before Vsx1 re-expression; arrow: the direction of lineage extension pointing toward older/early-born neurons. M: Medial; L: Lateral. Scale bar: 30μm. F’, Cross-sectional view of Vsx1 expression in the box-enclosed region in F. Magenta line encloses Grh-expressing cells (NE). Scale bar: 5μm. G, Cross-sectional view (as in B) of Bi expression relative to Grh-expressing neuroepithelial cells (magenta) and Dpn-expressing neuroblasts (yellow). Bi (cyan) is detected in the neuroepithelium and differentiated neurons, but not in the intervening progenitor zone. Asterisk: GMCs and nascent neurons before Bi re-expression; arrow: the direction of lineage extension pointing from younger/late-born toward older/early-born neurons. M: Medial; L: Lateral. Scale bar: 30μm. G’, Cross-sectional view of Bi expression in the box-enclosed region in G. Magenta line encloses Grh-expressing cells (NE). Scale bar: 5μm. NE, neuroepithelium; NB, neuroblast; GMC, ganglion mother cell.

Curiously, while Vsx1 and Bi are expressed in spatially restricted neuroepithelial domains in the outer proliferation center and later in defined subpopulations of neurons of the medulla cortex that emerge from these domains^16,17,19,28^ (**Figure 1E**), they are absent in NBs and in GMCs (**Figure 1F-G**). Despite this expression gap in progenitors, *Vsx1* is necessary for the determination of domain-specific neuronal fates^16^, implying that spatial information persists through progenitor stages. At the same time, some of the most abundant neurons in the medulla, such as Mi1 and Dm2, are specified independently of their domain of origin^15,16,27,29^. How spatial identity persists across progenitor stages in which spatial factors are absent, and how specific neurons escape it, remain unclear.

Here we show that an unexpectedly minimal mechanism resolves both problems. Spatial identity is encoded not through a transcriptional relay but as a precise and persistent chromatin signature restricted to a single spatial-factor locus: the *Vsx1/2* locus in Vsx progenies or the *Bi* locus in Bi progenies. This minimal trace enables neuronal re-expression of the same spatial factors to execute domain-specific differentiation programs. However, some neurons are able to escape this minimal specification and do not re-express spatial factors in a domain-specific fashion: They are specified independently of their spatial origin and produced from multiple domains. We further show that Polycomb repressive complex 2 (PRC2)-mediated repression restricts *Vsx1* re-expression to its home domain, while temporal identity and Notch-mediated fate selection determine which neurons engage or bypass the spatial program shortly after their terminal division. Thus, single-locus chromatin memory faithfully preserves spatial information while allowing specific neuronal fates to bypass it.

## Results

### Persistent single-locus chromatin accessibility distinguishes otherwise nearly indistinguishable domain progenies

While spatial factors are absent from NBs and GMCs, spatial identity persists through neurogenesis. To ask whether other domain-specific genes might bridge this gap or whether spatial information is instead retained at the chromatin level, we sorted NE cells and their progenies labeled by Gal4 reporters expressed in the Vsx, Optix, or Bi neuroepithelial domains: *pxb-T2A-Gal4* for the Vsx domain^27,30^, *Optix-Gal4* for the Optix domain^18,27^, and *dpp-Gal4* for the Bi domain^18,27^. Although these drivers are active primarily in neuroepithelial cells, perdurance of Gal4 and nuclear GFP maintains reporter signal in their NB, GMC, and young neuronal progeny, allowing us to recover these cells from each spatial origin (**Figure 2A**). We then profiled the sorted cells by single-cell multiome mRNA/ATAC-seq. To minimize confounding factors between spatial origin and sequencing batches, samples from the three reporter lines were pooled within each sequencing batch. Spatial origin was assigned to each cell by SNP demultiplexing using strain-specific variants from defined backgrounds in the *Drosophila* Genetic Reference Panel (DGRP)^26,31,32^ (**Extended Data Figure 1A**; Methods).

**Figure 2.**
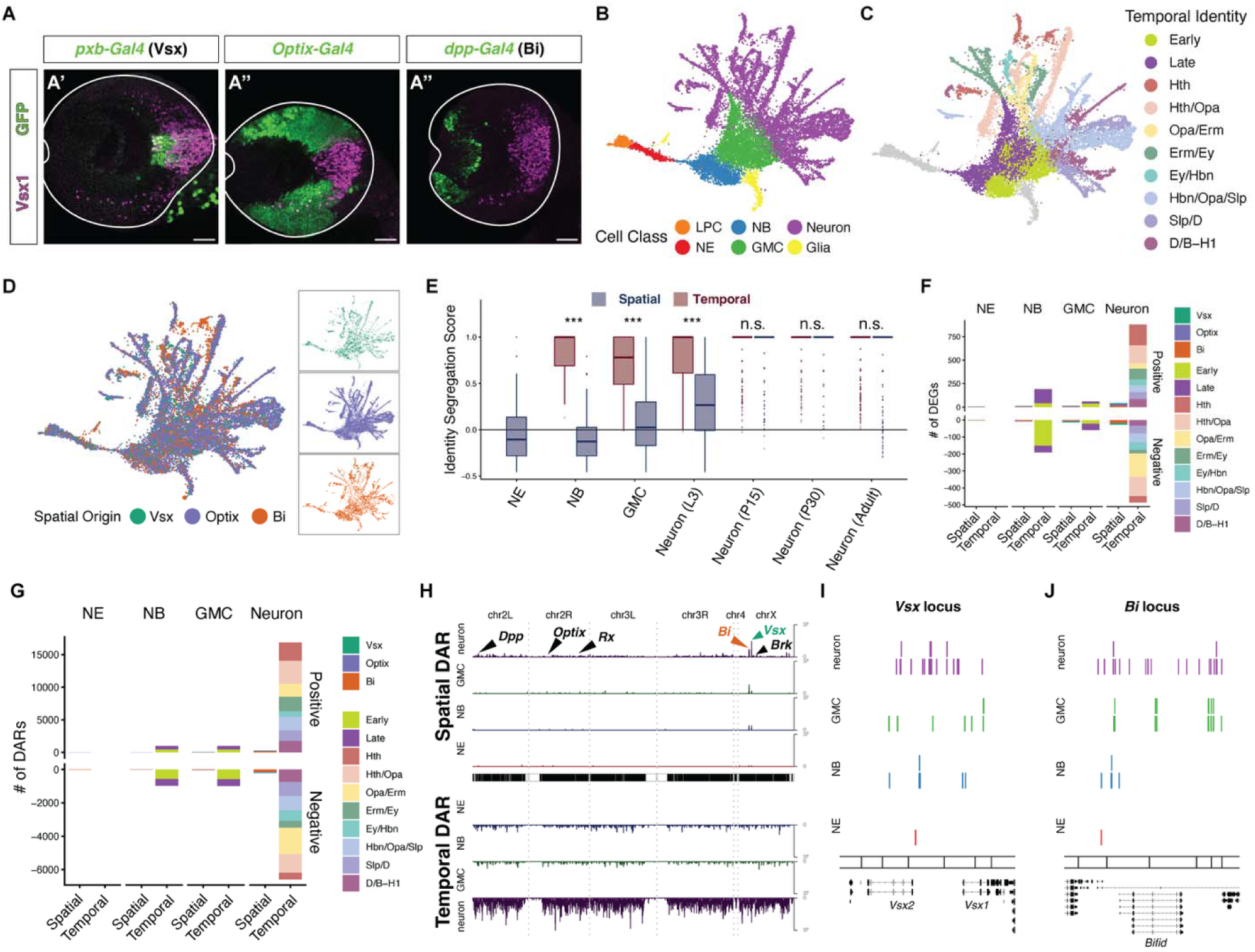
Single-cell multiome profiling reveals a precise domain-specific chromatin signature at spatial-factor loci despite minimal genome-wide divergence. A, Reporter expression used for spatial-origin profiling in third-instar larval optic lobes (A’: *pxb-T2A-Gal4*; A’’: *Optix-Gal4*; A’’’: *dpp-Gal4*). Green, nuclear GFP reporter; magenta, Vsx1, shown as a positional reference. Although these Gal4 drivers are active primarily in neuroepithelial cells, perdurance of Gal4 and nuclear GFP retains reporter signal in NB, GMC, and neuronal progenies, enabling recovery of cells by spatial origin. White outlines enclose the optic lobe. Scale bars: 30μm. B, Joint RNA-ATAC UMAP embedding of profiled cells colored by annotated cell class. LPCs were identified but excluded from downstream spatial comparisons. LPC, lamina precursor cell; NE, neuroepithelial cell; NB, neuroblast; GMC, ganglion mother cell. C, Same UMAP as in B, colored by temporal identity. Because individual temporal windows were incompletely resolved in NBs and GMCs, progenitor-stage temporal identities were grouped into early and late cohorts, while neuronal temporal identities were annotated at finer resolution with postmitotic temporal markers^22–24,33^. D, Same UMAP as in B, colored by spatial origin inferred from reporter genotype and SNP demultiplexing. Insets on the right show cells from each spatial origin plotted separately. E, Identity Segregation Score (ISS) comparing spatial origins (blue) and temporal identities (red) across developmental stages. Spatial ISS is near zero in NE, NB, and GMC populations and remains modest in third-instar neurons, while temporal ISS is high in NB, GMC, and neuronal populations. P15, P30, and adult neuron values are from reanalysis of published transcriptomic datasets, with spatial origin assigned by cell-type annotation rather than lineage tracking; types produced from multiple spatial domains were excluded from this analysis. Center line, median; box, interquartile range; whiskers, 1.5× interquartile range; ***P < 0.001; **P < 0.01; *P < 0.05; n.s., not significant; two-sided Wilcoxon signed-rank threshold test was performed to compare spatial and temporal ISS with a minimum difference threshold of 0.2. F, Number of differentially expressed genes (DEGs) identified by one-vs-rest pseudobulk comparisons within each cell class, separated by spatial origin or temporal identity. Positive and negative bars indicate genes enriched or depleted, respectively, in the indicated identity relative to the remaining identities in that comparison. Temporal comparisons in NB and GMC populations were performed using early and late cohorts. Genes with |log□FC| > 0.5, FDR < 0.05 were counted. G, Number of differentially accessible regions (DARs) identified by one-vs-rest pseudobulk comparisons within each cell class, plotted as in F. Regions with |log□FC| > 0.5, FDR < 0.05 were counted. H, Genome-wide ideogram-density plot of DARs across cell classes. Upper tracks show spatial DAR density; lower inverted tracks show temporal DAR density. Spatial DARs are concentrated at two spatial-factor loci, *Vsx1/2* and *Bi*, while temporal DARs are broadly distributed across the genome. Other spatial-patterning loci marked for reference, including *Dpp*, *Optix*, *Rx*, and *Brk*, show no comparable spatial DAR enrichment. I, Locus-level view of spatial DARs at the *Vsx1/2* locus. Each bar represents a unique DAR detected in the indicated cell class; DARs shared across multiple spatial-origin contrasts are collapsed by genomic coordinate. Across NE, NB, GMC, and neuronal stages, spatial DARs remain concentrated at this locus in Vsx-domain progenies. J, Locus-level view of spatial DARs at the *Bi* locus, plotted as in I. S Across NE, NB, GMC, and neuronal stages, spatial DARs remain concentrated at this locus in Bi-domain progenies. Gene models are shown below the DAR tracks in I and J.

After quality filtering and weighted-nearest-neighbor clustering (**Extended Data Figure 1B-D**), clusters were annotated as five broad cell classes using known marker genes: NEs, NBs, GMCs, neurons, and glia (**Figure 2B; Extended Data Figure 1E**). Temporal identity was assigned using known temporal TFs for NBs and GMCs and terminal selectors for neurons (**Figure 2C; Extended Data Figure 1F**).

Consistent with our prior work^22,29^, molecular diversity was primarily organized by temporal identity: neurons from different temporal windows were well-segregated in the joint RNA-ATAC UMAP embedding (**Figure 2C**). In contrast, cells from different spatial origins were extensively intermingled at every stage of neurogenesis, with each origin spanning broadly overlapping regions of the embedding when shown separately (**Figure 2D**). To quantify this asymmetry, we computed an Identity Segregation Score (ISS) for each cell, measuring the degree to which its local neighborhood is enriched for cells of the same identity relative to that identity’s global frequency (Methods). An ISS of zero indicates the mixing expected from global label frequencies, while values approaching one indicate nearly pure neighborhoods. Temporal ISS was high across NBs, GMCs, and neurons, confirming that temporal identity creates robust molecular distinctions once temporal patterning becomes active in NBs (**Figure 2E**). By contrast, although focused subclustering of NE cells alone can resolve spatial domains^22^, spatial ISS was near zero in NE, NB, and GMC populations and reached only modest levels in newborn neurons (**Figure 2E**), indicating little molecular segregation by spatial origin among progenitors and only modest segregation among newborn neurons. Re-analysis of spatially annotated pupal and adult neuronal types from an existing developmental atlas showed substantially higher spatial ISS in neurons^25,27^, suggesting that domain-associated molecular programs become more distinct during later neuronal maturation despite being weakly resolved in newborn neurons (**Figure 2E**; Supplementary Table 1; Methods).

Differential expression and accessibility analyses corroborated this asymmetry: across NBs, GMCs, and neurons, temporal identities accounted for far more differentially expressed genes (DEGs) and differentially accessible regions (DARs) than spatial origins (**Figure 2F-G**; Methods; Supplementary Tables 2-3). This disparity was not explained by the greater number of temporal identities tested, since the same asymmetry held when each identity was considered separately: at every stage, the spatial Bi domain that yielded the most DEGs and DARs still yielded many fewer than the temporal identity yielding the fewest (**Figure 2F-G**). For instance, the Bi-domain origin yielded 14 DEGs and 15 DARs in NBs compared with 191 DEGs and 974 DARs separating the early and late temporal cohorts. In neurons, Bi-domain origin again yielded fewer than Ey/Hbn, despite representing the most spatial and fewest temporal differences, respectively (32 versus 110 DEGs; 309 versus 1,492 DARs).

Despite their very small number, the spatial DARs were not distributed broadly across the genome but were concentrated at either of the two spatial-factor loci, *Vsx1/2* in cells from the Vsx domain and *Bi* in cells from the Bi domain. This concentration was observed in NE cells and persisted through NBs and GMCs (**Figure 2H**), stages at which neither the Vsx1 nor the Bi protein was detectable (**Figure 1F-G**). *Vsx1* and *Bi* themselves were also among the spatial DEGs in both NBs and GMCs. HCR detected only transcription-site puncta at both loci in *Mira*-positive neuroblasts without cytoplasmic mRNA accumulation (**Extended Data Figure 1G-H**). Examination of the *Vsx1/2* and *Bi* loci revealed multiple differentially accessible regions not only within the transcribed gene bodies but also at flanking regulatory elements of either locus.

Although most individual spatial DARs were not maintained across cell classes, domain-specific accessibility remained concentrated at the *Vsx1/2* locus in Vsx-domain progenies and at the *Bi* locus in Bi-domain progenies (**Figure 2I-J**). In contrast, no spatial DARs were detected at the *Optix* locus at any stage, even in neuroepithelial cells, where *Optix* is expressed (**Extended Data Figure 1J**). *Optix* is not re-expressed in neurons emerging from its spatial domain. Optix-domain progenies therefore carry no persistent chromatin trace of their spatial origin. Other spatial-patterning loci, including *Dpp*^15,19^, *Rx*^16^, and *Brk*^15^, likewise showed neither comparable spatial-DAR enrichment nor persistent progenitor-stage differential accessibility (**Figure 2H**; **Extended Data Figure 1I-L**).

Together, these results show that progenies from different spatial domains remain nearly indistinguishable in transcriptome and chromatin accessibility while retaining a precise, locus-specific chromatin signature at either the *Vsx1/2* or the *Bi* locus.

### Neuronal re-expression of spatial factors specifies domain-specific fates

Although Vsx1 and Bi proteins are not detectably expressed in NBs or GMCs, both factors reappear in subsets of postmitotic medulla neurons produced from their respective spatial domains (**Figure 1E-G**)^17,25,27^. Together with the persistence of domain-specific chromatin accessibility at the *Vsx1/2* or *Bi* loci through the intervening progenitor stages (**Figure 2H-J**), this suggests that neuronal re-expression executes spatial identity. To test this, we used a temperature-sensitive Gal80 repressor to restrict the activity of a pan-neuronal Gal4 driver (*elav^C155^-Gal4*) to the final 48 hours of larval development, when medulla neurons are generated from the outer proliferation center (**Extended Data Figure 2A**; Methods).

We first tested whether *Vsx1/2* is required for Vsx-domain neuronal fates across two temporal cohorts. In the early-born Run^+^ temporal cohort, control animals generate both Pm4 neurons (Ct^−^ Run^+^ from the Vsx domain) and Mi4 neurons (Ct^+^ Run^+^ from the Optix domain)^22,27^. Pan-neuronal knockdown of *Vsx1* and *Vsx2* resulted in Mi4 neurons, normally confined to the Optix domain, appearing throughout the anterior Vsx territory at the expense of Pm4 neurons (**Figure 3A-C**; ∼79% reduction in Pm4; p = 0.037). Knockdown of either *Vsx* paralog alone produced no detectable change for Pm4 (**Extended Data Figure 2B-D**), consistent with functional redundancy between *Vsx1* and *Vsx2*^34^. In a later-born temporal cohort marked by *Ey* and *Kn* co-expression^22,29,35,36^, Vsx1/2 loss of function similarly reduced Tm5e, the Vsx-domain progeny of this cohort (*Ey^+^ Kn^+^ D^-^*), relative to its *Ey^+^ Kn^+^ D^+^* Optix-domain alternatives, *e.g*. TmY5a (**Figure 3D-F**; ∼77% reduction in anterior Tm5e; p = 0.0025). Thus, *Vsx1/2* is required for Vsx-domain fate selection across the two temporal windows examined.

**Figure 3.**
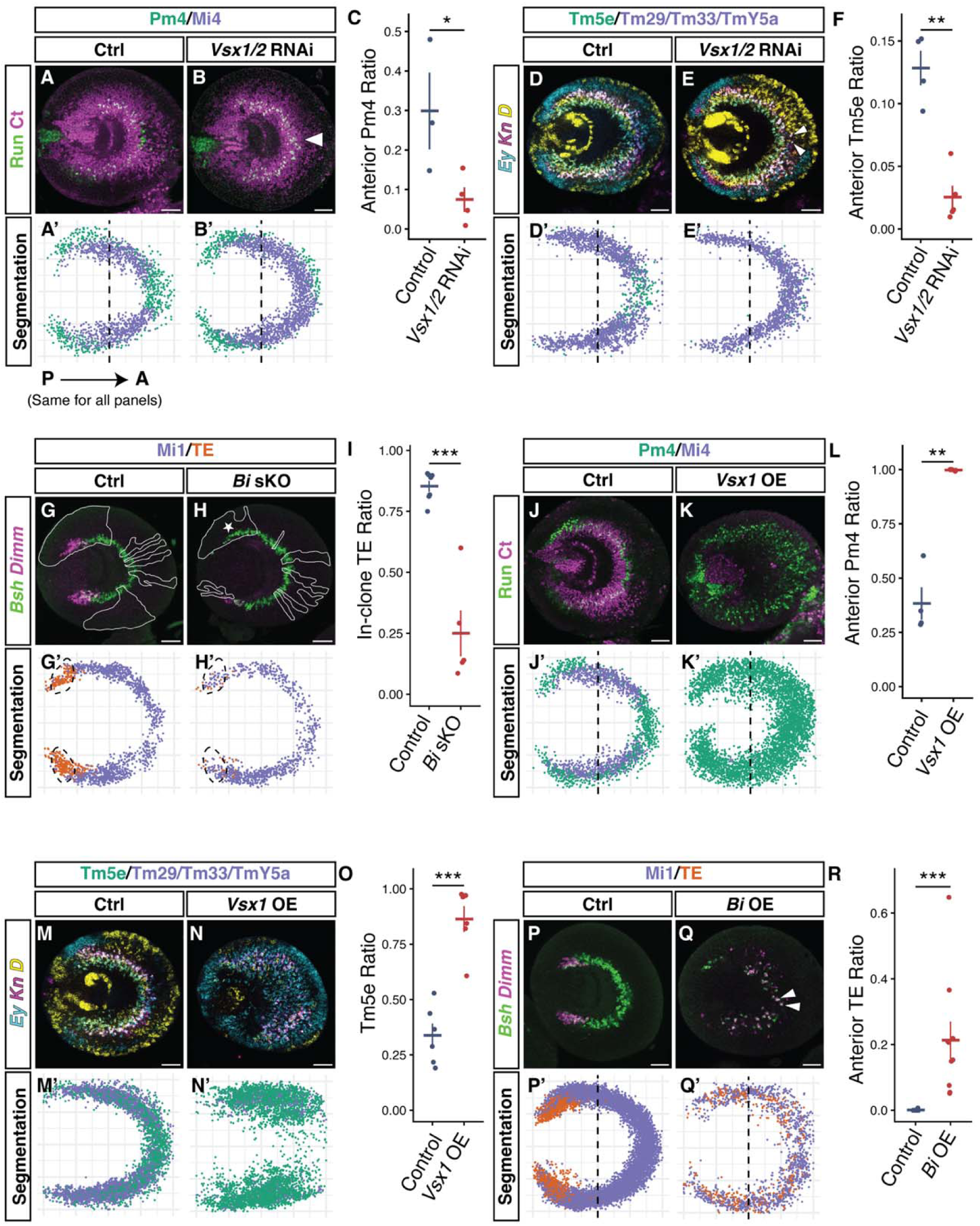
Vsx1/2 and Bi perturbations shift domain-specific neuronal fates. For each perturbation, representative immunostaining (A-B and J-K) or HCR-FISH images (D-E, G-H, M-N, and P-Q) are shown for the control and perturbation conditions. Prime panels show pooled segmentation overlays from all quantified brains for the temporal cohort or neuronal fate of interest. Each segmented cell is projected into a common coordinate system and colored by marker-defined domain-associated identity. All panels are oriented so that the right is anterior. The dashed lines in A’, B’, D’, E’, J’, K’, P’ and Q’ mark the antero-posterior boundary. Quantification panels show the corresponding fate proportion. Dot plots show biological replicates, with each point representing one larva; crossbars indicate mean ± SEM. Statistical significance was assessed using quasibinomial generalized linear models across larvae. * p < 0.05, ** p < 0.01, *** p < 0.001. Scale bars: 30μm. A-C, Neuronal *Vsx1/2* loss of function shifts the early-born Run+ cohort from Pm4 toward Mi4 identity. A, A’, Control optic lobe (mCherry knockdown) stained for Run and Ct, with segmentation of Run+ cells colored as Pm4 (Ct− Run+) or Mi4 (Ct+ Run+). B, B’, *Vsx1/2* knockdown. White arrowhead indicates a region exclusively Run+/Ct− in controls, now occupied by Run+/Ct+ cells. C, Quantification of Pm4 proportion among anterior Run+ cells; n = 3 control and 4 *Vsx1/2* knockdown larvae. D-F, Neuronal *Vsx1/2* loss of function similarly shifts a later-born *Ey+ Kn+* cohort away from Tm5e identity. D, D’, Control optic lobe (mCherry knockdown) stained for *Ey, Kn*, and *D*, with segmentation of *Ey+ Kn+* cells colored as Tm5e (D−) or Optix-domain alternative fates (D+; including Tm29, Tm33, and TmY5a). E, E’, *Vsx1/2* knockdown. White arrowheads indicate *Ey+ Kn+ D+* cells in the anterior domain. F, Quantification of Tm5e proportion among anterior *Ey+ Kn+* cells; n = 4 control and 5 *Vsx1/2* knockdown larvae. G-I, *Bi* loss of function reduces TE neurons. G, G’, Control optic lobe (Cas9 only) showing TE neurons marked by *Dimm* and *Bsh*, with segmentation of *Bsh+* cells colored as TE or Mi1. H, H’, Somatic *Bi* CRISPR. Star indicates a large clone covering the TE territory. I, Quantification of TE proportion (*Dimm+ Bsh+*) among *Bsh+* cells within clones, in the dashed-line-enclosed TE territory defined by kernel density estimation on control samples (Methods); n = 7 control and 5 *Bi* CRISPR larvae. J-L, Neuronal *Vsx1* overexpression is sufficient to promote Pm4 identity in the Run+ cohort. J, J’, Control optic lobe (*LacZ* overexpression) stained for Run and Ct, with segmentation of Run+ cells colored as Pm4 or Mi4. K, K’, *Vsx1* overexpression. L, Quantification of Pm4 proportion among anterior Run+ cells; n = 4 control and 4 *Vsx1* overexpression larvae. M-O, Neuronal *Vsx1* overexpression is sufficient to promote Tm5e identity in the *Ey+ Kn+* cohort. M, M’, Control optic lobe (*LacZ* overexpression) stained for *Ey, Kn*, and *D*, with segmentation of Ey+ Kn+ cells colored as Tm5e or *D+* Optix-domain alternatives. N, N’, *Vsx1* overexpression. O, Quantification of Tm5e proportion among *Ey+ Kn+* cells; n = 6 control and 6 *Vsx1* overexpression larvae. P-R, Neuronal *Bi* overexpression is sufficient to promote TE fate. P, P’, Control optic lobe (*LacZ* overexpression) showing TE neurons marked by *Dimm* and *Bsh*, with segmentation of anterior TE cells. Control cells scored as anterior lie immediately adjacent to the domain boundary and reflect normalized positions falling marginally across x = 0. Q, Q’, *Bi* overexpression. White arrowheads indicate *Dimm+/Bsh+* cells in the anterior domain. R, Quantification of anterior TE proportion among *Bsh+* cells; n = 9 control and 10 *Bi* overexpression larvae.

We then tested whether *Bi* is required for a Bi-associated neuronal fate. Because multiple independent RNAi lines failed to knock down *Bi* in the optic lobe (**Extended Data Figure 2E-G**), we instead pursued somatic disruption of *Bi* using CRISPR. In the earliest (Hth) temporal cohort, the Bi domain produces a subclass of TE neurons (*Bsh^+^ Dimm^+^*), while the Vsx and Optix domains produce Mi1 (*Bsh^+^ Dimm^−^*). In the mutant, there was a consistent loss of TE neurons that appeared to be transformed into Mi1-like identity (**Figure 3G-I and Extended Data Figure 2H-M**; ∼81% reduction in TE; p = 4.4 × 10^-6^). In this experiment, Cas9 expression was induced by heat shock-activated FLP recombinase three days before dissection, excising a transcriptional stop cassette under the ubiquitous *Act5C* promoter (Methods). Because CRISPR was induced early and was not neuron-restricted, whether *Bi* is required in the neuroepithelium, in neurons, or in both remains unresolved. Nevertheless, these data establish that *Bi* function during development is required for TE specification.

We next asked whether neuronal expression of spatial factors is sufficient to redirect cell fates. Pan-neuronal overexpression of *Vsx1* was sufficient to drive Vsx-domain fates in both Vsx-Optix fate pairs tested above. In the Run^+^ cohort, pan-neuronal *Vsx1* overexpression nearly eliminated Mi4 identity, and Pm4 consequently accounted for nearly all anterior Run^+^ neurons (**Figure 3J-L**; anterior Mi4 proportion reduced from ∼39% to ∼0.4%; p = 0.0021). In the *Ey^+^ Kn^+^* cohort, *Vsx1* overexpression in neurons similarly increased Tm5e while depleting the *D^+^* Optix-domain alternatives (**Figure 3M-O**; Optix-domain fates reduced from ∼68% to ∼18%; p = 8.8 × 10^-4^). Pan-neuronal overexpression of *Bi* likewise increased TE abundance across the optic lobe and induced TE neurons ectopically throughout the anterior Vsx and Optix progenies in the medulla, a territory in which TE neurons are absent in controls (**Figure 3P-R**; **Extended Data Figure 2N**; anterior TE proportion increased from ∼0.1% to ∼21%; p = 6.3 × 10^-4^).

Therefore, both loss- and gain-of-function perturbations produced reciprocal changes in domain-specific neuronal fates. *Vsx1/2* loss converted two Vsx-domain fates toward Optix-domain alternatives while ectopic *Vsx1* drove the corresponding Vsx-domain identities at the expense of Optix-domain alternatives. *Bi* loss depleted TE neurons, while neuronal *Bi* overexpression induced TE ectopically. These reciprocal perturbations also reveal a difference in how spatial identity is executed in the Vsx and Bi domains *vs.* the Optix domain. *Optix* itself is turned off in NB and GMCs and it is not re-expressed in medulla neurons. Although *Optix* perturbation during neuroepithelial patterning alters neuronal fate^16^, we did not identify terminal selectors shared broadly among Optix-domain progenies^27^. Although the Optix domain is further subdivided by additional spatial programs that can generate distinct neuronal fates^15,30^, the Optix-domain neural types examined here are produced across these subdivisions^27^. Therefore, Optix-domain fates like Mi4 and TmY5a appear to be neurons specified according to their temporal cohort, while *Vsx1/2* and *Bi* modify these temporal outputs by regulating specific postmitotic marker combinations: *Vsx1/2* suppresses *Ct* in the Run^+^ cohort (**Figure 3A-C, J-L**) and *D* in the *Ey^+^ Kn^+^* cohort (**Figure 3D-F, M-O**), whereas *Bi* activates *Dimm* in the *Bsh^+^* cohort (**Figure 3G-I, P-R**). This suggests that these Optix-domain fates represent a default-like state of the temporal program, from which *Vsx1/2* or *Bi* re-expression diverts neurons into domain-specific alternatives.

Together, these results support a model in which neuronal re-expression of spatial factors executes domain-specific differentiation programs, while neurons that do not engage these programs ignore spatial domains and adopt Optix-like identity (see below)^15,16,27,37^. The necessity and sufficiency of *Vsx1/2* and *Bi* for domain-specific fates raise two related questions: how is their re-expression restricted to the correct domain, and how do neuron types that are generated across multiple domains adopt equivalent fates regardless of origin? We first addressed the mechanism of domain restriction.

### Polycomb restricts Vsx1 re-expression to its home domain

Domain-specific accessibility at the *Vsx1/2* and *Bi* loci persists through progenitor stages despite turnover of the specific differentially accessible elements across cell classes (**Figure 2H-J**), suggesting that home-domain progenies maintain a locus-level permissive state that could support later re-expression. However, not all prior neuroepithelial expression of spatial markers invariably leads to neuronal re-expression, as illustrated by *Optix*, which is not re-expressed in medulla neurons. Moreover, we did not detect additional domain-enriched transcriptional regulators in NBs or GMCs that could provide a lineage-intrinsic relay of spatial activating input into neurons (Supplementary Table 2). The lack of an apparent regulator relay prompted us to examine whether the *Vsx1/2* and *Bi* loci have a broad intrinsic capacity for activation, with domain-specific re-expression arising because the loci are repressed outside their home domains.

One candidate source of spatial repression is the cross-regulatory network that initially establishes the Vsx-Optix boundary in the OPC. However, this relationship is asymmetric: loss of *Vsx1* derepresses *Optix* in the neuroepithelium, while *Optix* is sufficient to repress *Vsx1* when mis-expressed but is not required to exclude it from the Optix domain^16,38^. Moreover, neither factor is expressed in NBs or GMCs, and *Optix* itself is not expressed in medulla neurons. Thus, the known *Vsx1*-*Optix* cross-repression cannot account for the continued spatial restriction of neuronal *Vsx1* re-expression.

Like many developmental regulator loci, the *Vsx1/2* and *Bi* loci carry H3K27me3 marks in the brain at the third larval instar, as demonstrated by bulk CUT&RUN profiling^39^. Consistently, targeted DamID detects persistent Polycomb (Pc) protein binding at these loci through neuroblast, ganglion mother cell, and neuronal stages^40^ (**Extended Data Figure 3A-B**). These observations raised the possibility that the PRC pathway provides the active repressive layer posited above.

To test whether PRC2 is required to restrict *Vsx1* re-expression, we knocked down the PRC2 catalytic subunit *E(z)* specifically in neurons using the same pan-neuronal driver with a temperature-sensitive Gal80 repressor that was used for spatial factor perturbations (**Extended Data Figure 2A**; Methods). In control animals, Vsx1 expression was largely restricted to neurons in the anterior medulla cortex generated by the Vsx neuroepithelial domain, with rare posterior expression corresponding to domain-independent Vsx1-expressing types^15,29,34^ (**Figure 4A-A’**; see next section). However, upon neuronal *E(z)* knockdown, Vsx1 expression expanded into neurons of the posterior medulla cortex, which contains Optix- and Bi-domain progenies (**Figure 4B-B’**). Among Vsx1-positive neurons, the proportion located in the posterior medulla rose from 3.6% in controls to 13.0% after *E(z)* RNAi (p = 0.00055; **Figure 4C**). Ectopic Vsx1 expression was biased toward later-born neurons, with most de-repression occurring after the Ey temporal window (Svp^+^)^22^ (**Extended Data Figure 3E-F**). This expansion is consistent with ectopic engagement of the normally domain-restricted *Vsx1* re-expression program. However, *E(z)* knockdown did not affect the fraction of early-born Notch^ON^ neurons that are domain-ignoring types and normally do not express Vsx1 (See below and **Extended Data Figure 3G**). Thus, loss of *E(z)* allows *Vsx1* re-expression outside the Vsx domain but does not induce *Vsx1* in Vsx-domain neurons that normally lack it.

**Figure 4.**
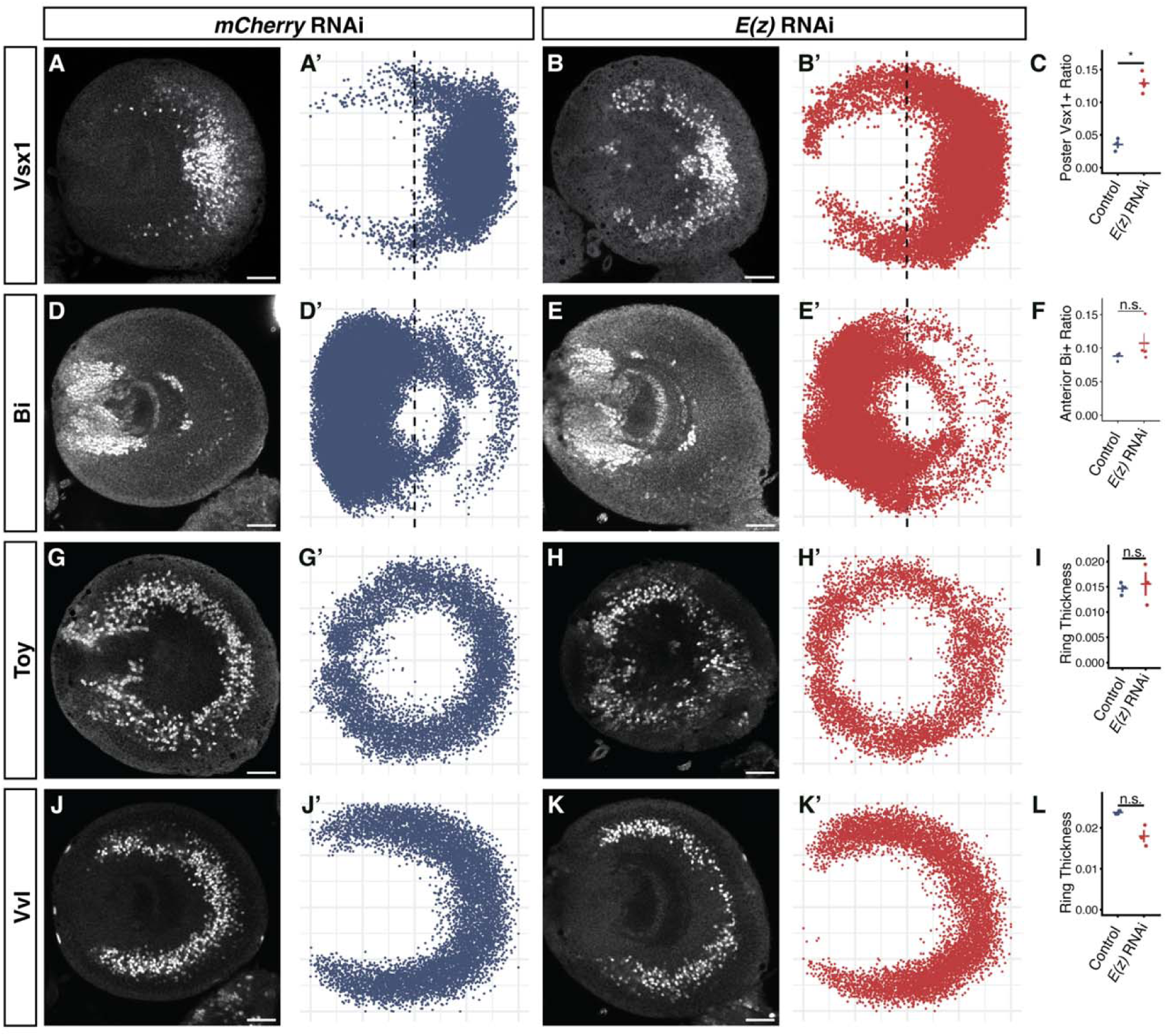
PRC2 selectively restricts Vsx1 re-expression to its spatial domain. For each perturbation, representative immunostaining images are shown for the control and perturbation conditions. Prime panels show pooled segmentation overlays from all quantified brains for the gene of interest. Each segmented cell is projected into a common coordinate system. A-C, Neuronal *E(z)* knockdown causes ectopic Vsx1 expression outside the Vsx domain. A, A’, Vsx1 expression and segmentation overlay in control optic lobes (*mCherry* RNAi; n = 3 larvae). B, B’, Vsx1 expression and segmentation overlay in optic lobes expressing *E(z)* RNAi (n = 4 larvae). C, Quantification of the fraction of all Vsx1-positive neurons located in the posterior half of the medulla cortex. D-F, Neuronal *E(z)* knockdown does not significantly expand Bi expression. D, D’, Bi expression and segmentation overlay in control optic lobes (*mCherry* RNAi; n = 4 larvae). E, E’, Bi expression and segmentation overlay in optic lobes expressing *E(z)* RNAi (n = 4 larvae). F, Quantification of the fraction of all Bi-positive neurons located in the anterior half of the medulla cortex. The dashed lines in A’, B’, D’ and E’ mark the antero-posterior boundary. Bi-expressing domain size varies with medio-lateral depth, with deeper layers being wider. The wider domain is reflected in projected overlays D’ and E’. G-I, Neuronal *E(z)* knockdown does not alter the temporal distribution of Toy expression. G, G’, Toy expression and segmentation overlay in control optic lobes (*mCherry* RNAi; n = 3 larvae). H, H’, Toy expression and segmentation overlay in optic lobes expressing *E(z)* RNAi (n = 3 larvae). I, Variance of the radial positions (ring thickness) of Toy-positive neurons. The radial axis of the medulla cortex corresponds to neuronal birth order, with early-born neurons closer to the center; radial variance was therefore used as a proxy for the breadth of the birth-order range over which Toy is expressed. J-L, Neuronal *E(z)* knockdown does not expand the temporal distribution of Vvl expression. J, J’, Vvl expression and segmentation overlay in control optic lobes (*mCherry* RNAi; n = 4 larvae). K, K’, Vvl expression and segmentation overlay in optic lobes expressing *E(z)* RNAi (n = 3 larvae). L, Variance of the radial positions (ring thickness) of Vvl-positive neurons, quantified as in I. The observed difference was toward reduced radial variance, rather than expansion of the Vvl expression domain. Quantification plots show individual biological replicates, with each point representing one larva; crossbars indicate mean ± SEM. Statistical significance in C and F was assessed across larvae using quasibinomial regression, whereas radial variances in I and L were compared using two-sided Welch’s t-tests. *P < 0.05; n.s., P ≥ 0.05. Scale bars: 30μm.

To determine whether this reflected selective sensitivity of the *Vsx1* locus or broad de-repression of H3K27me3-marked developmental regulators, we examined *Bi* and the temporal terminal selectors *Toy* and *Vvl*, whose loci likewise carry H3K27me3 and persistent Polycomb occupancy. As for Vsx, Bi is absent in NBs and GMCs but *Bi* gets re-expressed in neurons. However, the proportion of Bi-positive neurons located in the anterior medulla cortex did not change significantly after *E(z)* knockdown: Approximately 8.8% in controls (including Bi-domain-specific Lawf neurons that had migrated anteriorly^41,42^) *vs*. 10.5% in *E(z)* RNAi; p = 0.217 (**Figure 4D-F**; **Extended Data Figure 3B**). Toy and Vvl provide a complementary test because their expression is restricted along the temporal rather than spatial axis: Successive temporal cohorts of medulla neurons occupy successive concentric positions. Thus, the thickness of a ring represents the number of temporal windows in which a gene is expressed, and loss of temporal restriction would therefore expand the thickness of these rings rather than shift them across domain boundaries (**Figure 4G and J**). Following neuronal *E(z)* knockdown, neither expression domain expanded; Toy ring thickness was unchanged (p = 0.74), whereas Vvl showed a nonsignificant trend toward contraction rather than expansion (p = 0.06; **Figure 4G-L**; **Extended Data Figure 3C-D**).

This selective sensitivity to *E(z)* loss suggests that *Vsx1* differs from other H3K27me3-marked regulators not simply in its repressive state, but in the presence of neuronal activating input whose readout must normally be restricted to the Vsx domain by PRC2 acting at the endogenous locus. To test whether the *Vsx1/2* locus contains such broadly responsive neuronal regulatory activity, we generated reporters from domain-enriched accessible regions upstream of the *Vsx1* transcription start site, including *Fragment 1* (2.1 kb, approximately 6.3 kb upstream), previously shown to be active in Vsx1-expressing NE cells^38^, and *Fragments 2* and 3 (1.2 and 3.1 kb, approximately 10.3 and 14.9 kb upstream), recently identified as differentially accessible in Vsx-domain neurons^29^ (**Extended Data Figure 3H-K**). A Gal4 reporter based on *Vsx-Fragment 1* drove domain-restricted GFP reporter expression in *Vsx1*-expressing NEs and their progenies, but Gal4 transcripts were not detected in NBs, GMCs, and neurons, indicating that its neuronal GFP signal reflected perdurance (**Extended Data Figure 3L-O**). In contrast, Gal4 reporters based on *Vsx-Fragments 2 and 3* showed no detectable progenitor activity but drove broad neuronal GFP reporter expression around the medulla crescent without obvious domain preference (**Extended Data Figure 3P-Q’**).

Thus, isolated neuronal regulatory elements from the *Vsx1/2* locus can respond to activating inputs outside the Vsx domain when removed from their endogenous locus context. Broad neuronal activity from isolated fragments could reflect nonspecific out-of-context expression rather than genuine engagement of Vsx1-activating inputs; however, within the Vsx territory, reporter expression from both fragments was almost perfectly concordant with endogenous Vsx1 expression (*Fragment 2*: 99.0 ± 0.05%, n = 3; *Fragment 3*: 100%, n = 3; **Extended Data Figure 3R**). This supports the interpretation that these elements capture neuronal regulatory activity used by the endogenous *Vsx1/2* locus. At the endogenous locus, locus-level repression, including PRC2-mediated silencing, contributes to restricting this activity to the Vsx domain.

Together, these results support a model in which the Vsx1/2 locus contains neuronal regulatory elements with broad intrinsic activity, while PRC2-mediated silencing at the locus confines their activity to Vsx-domain progenies. This may reflect domain-specific states of the same locus: persistent transcriptional activity could resist PRC2-mediated silencing^43,44^, while the same locus in non-home progenies lacks such protection and remains silenced. This PRC2-dependent restriction, however, is not generalizable to *Bi*, whose re-expression remained domain-restricted after *E(z)* knockdown, suggesting that spatial restriction is achieved by more than one mechanism.

### Temporal and Notch identity control domain specificity of neuronal *Vsx1/2* expression

Having established how PRC2 maintains the domain restriction of *Vsx1* re-expression, we next asked how some neurons become independent of their domain of origin. Several medulla neuron types are generated from both the Vsx and Optix domains and adopt equivalent fates regardless of origin^15,16,27,37^ (**Extended Data Figure 4A**). We refer to these as domain-ignoring types with respect to the Vsx-Optix distinction. If *Vsx1/2* re-expression executes Vsx-domain identity, domain-ignoring neurons could achieve uniformity in two ways: by failing to re-express *Vsx1/2* even when born from the Vsx domain, or by activating *Vsx1/2* through a domain-independent regulatory program when born elsewhere. Both strategies appear to be used.

Multiple domain-ignoring neuronal types, including Mi1 and Tm1/2/4/6, lack *Vsx1* expression at the third larval instar even though a proportion of each of these neuronal types originate from the Vsx domain (**Figure 5A-B’**). In third-instar neuronal ATAC-seq profiles pooled by temporal and Notch identity, these domain-ignoring neurons showed broadly reduced accessibility across the *Vsx1/2* locus relative to Vsx-domain neurons that re-express *Vsx1/2,* e.g. Pm3 or Pm4 (**Figure 5C; Extended Data Figure 4C-E**). This included lower accessibility in the regions of Vsx-Fragments 2 and 3, which were instead accessible and exhibited broad neuronal reporter activity in Vsx-domain neurons that re-expressed Vsx (**Extended Data Figure 3P-Q’**). Reduced accessibility at these domain-bookmarked regulatory elements is consistent with these fates being specified without activating the Vsx-domain re-expression program.

**Figure 5.**
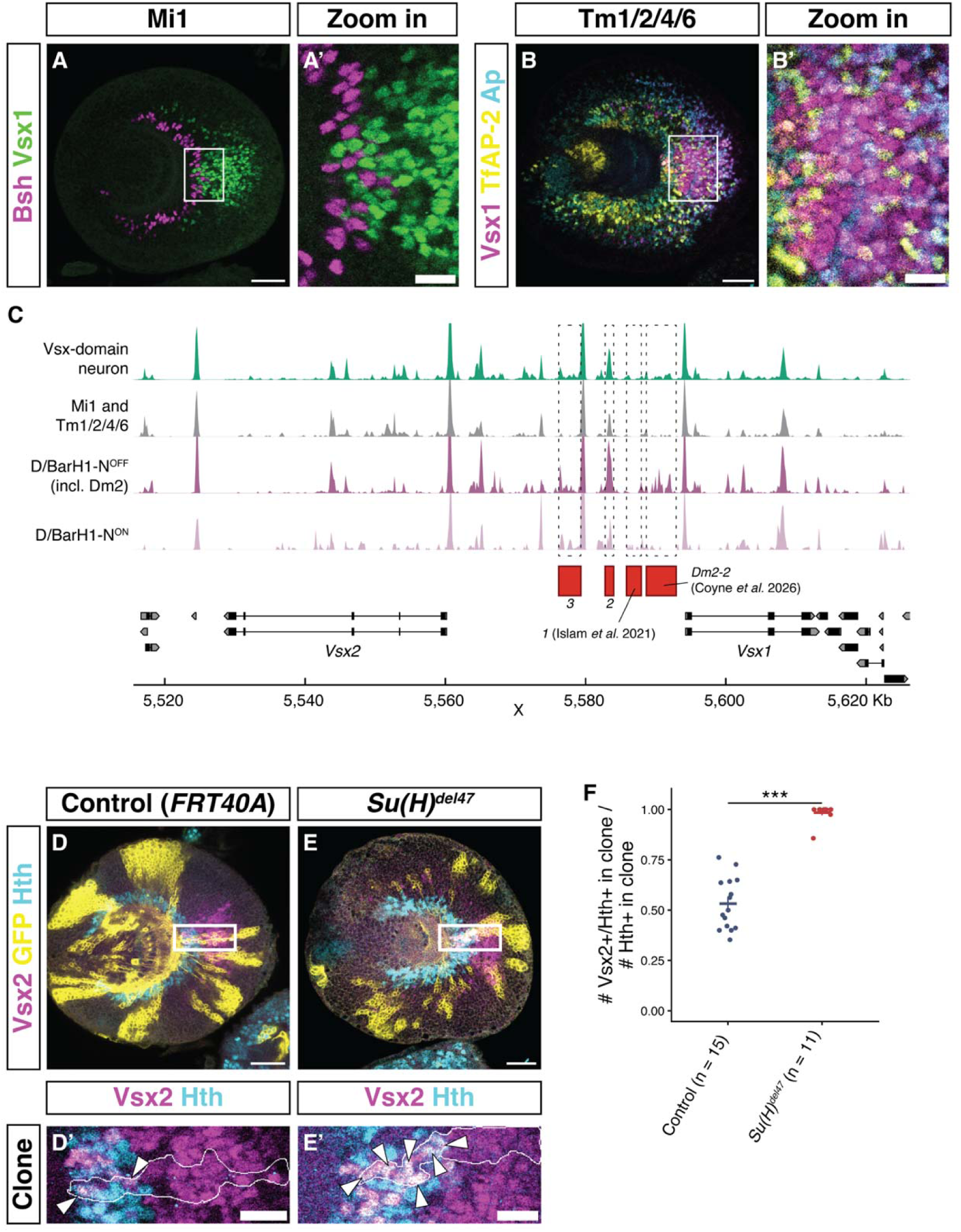
Notch-dependent fate identity determines engagement with the Vsx1/2 re-expression program. A-B, Early-born domain-ignoring neuron types lack Vsx1 expression. A, A’, Optic lobe stained for Bsh (magenta) and Vsx1 (green). A’, Zoom of the anterior medulla (boxed region in A) showing that Bsh-positive Mi1 neurons do not co-express Vsx1. B, B’, Optic lobe stained for Vsx1 (magenta), TfAP-2 (yellow), and Ap (cyan). Tm1/2/4/6 neurons are marked by TfAP-2 and Ap co-expression. B’, Zoom of boxed region in B showing that TfAP-2+/Ap+ neurons do not co-express Vsx1. C, Pseudobulk ATAC-seq accessibility across the Vsx1/2 locus in third-instar neuronal subsets. From top to bottom: Vsx-domain neurons excluding Mi1 and Tm1/2/4/6 (teal), Mi1 and Tm1/2/4/6 (gray), Vsx-domain D/BarH1 Notch^OFF^ cohort including Dm2 (dark purple), Vsx-domain D/BarH1 Notch^ON^ cohort (light purple). Mi1 and Tm1/2/4/6 neurons show broadly reduced accessibility across the locus relative to Vsx-domain neurons, while the D/BarH1 Notch^OFF^ cohort shows selective accessibility at the Dm2-2 element. Red bars below tracks indicate the positions of regulatory *Fragments 1, 2*, and *3* (also see Extended Data Figure 3H-K and 4C-F) and the *Dm2-2* element identified by Coyne et al.^29^. Gene models for Vsx2 and Vsx1 are shown at bottom. D-F, Loss of Notch signaling converts Mi1 to a Vsx2-positive Pm3 fate in the Vsx domain (see also Extended Data Figure 4G-H). Vsx2 was used as a proxy for *Vsx1/2* re-expression because Vsx1 and Vsx2 are co-expressed in Vsx-domain neurons (Extended Data Figure 4B), and Vsx1 could not be co-stained with Hth because the two antibodies were raised in the same host species. D, D’, Control MARCM clone (*FRT40A*) in the Vsx domain stained for Vsx2 (magenta), GFP (yellow), and Hth (cyan). D’, Zoom of the boxed region in D; clone boundary outlined in white; arrowheads indicate Vsx2+/Hth+ neurons within the clone. E, E’, *Su(H)^del47^* MARCM clone in the Vsx domain, displayed as in D, D’. Arrowheads indicate Vsx2+/Hth+ neurons. F, Fraction of Vsx2+/Hth+ neurons among all Hth+ neurons within Vsx-domain clones. Clones from the same animal were pooled before quantification; each dot represents one animal (n = 15 control animals; n = 11 *Su(H)^del47^*animals). Crossbars indicate mean ± SEM. Statistical significance assessed by quasibinomial regression. ***P < 0.001. Scale bars: 30μm.

In the medulla, Notch-mediated binary fate choice at the terminal GMC division generates molecularly distinct siblings within each temporal cohort^20,21,45^, and domain-ignoring types are typically born with a domain-specific sister cell (**Extended Data Figure 4A**). For example, Mi1 (Notch^ON^), which ignores its domain of origin, is the sibling of Pm3 (Notch^OFF^), which originates from the Vsx domain and re-expresses Vsx1/2^16^. Similarly, Notch^ON^ Tm1/Tm4 and Tm2/Tm6 ignore their domain of origin while their Notch^OFF^ siblings (*e.g.* for Tm1/Tm4, likely Pm4 in the Vsx and Mi4 in the Optix domain)^22,27^ do respect it. However, in other temporal windows, it is the Notch^OFF^ neuron (*e.g.* T1 neuron) that is domain ignoring. Two sisters can also often be both domain specific. However, when a cell is domain-ignoring its sister is in most cases domain-specific (**Extended Data Figure 4A**). This suggests that this binary choice also determines whether a neuron engages the spatial re-expression program. To test this, we generated Vsx-domain MARCM clones homozygous for the *Suppressor of Hairless* loss-of-function allele *Su(H)^del47^*, thereby removing the transcriptional effector of Notch signaling^20^, and examined Vsx2 expression as a proxy for *Vsx1/2* re-expression among Hth-positive neurons (Mi1 and Pm3) within clones (Vsx1 and Vsx2 are co-expressed in neurons; **Extended Data Figure 4B**).

In wild-type clones, approximately half of Hth^+^ neurons expressed Vsx2, consistent with each GMC division producing one Vsx2-positive Notch^OFF^ daughter (Pm3, marked by Svp) and one Vsx2-negative Notch^ON^ daughter (Mi1, marked by Bsh) (**Figure 5D-D’**). In *Su(H)^del47^*mutant clones, in which all daughters adopt Notch^OFF^ fates^20,21^, virtually all Hth^+^ neurons expressed Vsx2 (∼50% in wild-type vs. ∼100% in *Su(H)^del47^*; p = 1.85 × 10^-5^; **Figure 5E-F**). Consistent with fate conversion, Mi1 neurons were replaced by Pm3 neurons in mutant clones (**Extended Data Figure 4G-H**). Because Notch determines which sibling fate is specified, differences between siblings in *Vsx1/2* re-expression are assigned by that choice, though often both siblings re-express *Vsx1/2*. This fate conversion establishes that the domain-restricted chromatin trace is necessary but not sufficient for *Vsx1/2* re-expression: within the Vsx domain, whether a neuron engages the re-expression program depends on its Notch and temporal status.

However, domain independence does not always require excluding *Vsx1*. For example, Dm2 and Tm5c that are produced from both Vsx and Optix domains express *Vsx1* regardless of spatial origin^15,27,34^. Neuroepithelial *Vsx1* knockdown diminishes Vsx1 expression in domain-specific neurons but spares its expression in these domain-independent neurons (**Extended Data Figure 4I-L**), including when Dm2 and Tm5c emerge from the Vsx domain, indicating that domain-independent *Vsx1* expression does not depend on the neuroepithelial program. Therefore, *Vsx1* is controlled by at least two regulatory programs, one bookmarked by Vsx in its expression domain, and one that is independent from NE Vsx. Coyne and colleagues identified a *Vsx1/2* regulatory element, *Dm2-2*, and showed that it exhibits Dm2-enriched chromatin accessibility in both Vsx and Optix domains.

and that an activity reporter for this element is active in Dm2 neurons spanning all spatial domains^29^. Within the late D/BarH1 temporal window, the Notch^OFF^ cohort containing Dm2 showed accessibility at the *Dm2-2* element that was diminished in its Vsx domain specific sister and in other Vsx-domain neurons (**Figure 5C**; **Extended Data Figure 4F**). These observations are consistent with some neuronal fates expressing *Vsx1* through fate-specific regulatory elements independently of the domain-bookmarked program.

Notch status by itself, however, does not predict whether a neuronal fate expresses *Vsx1*. Vsx-domain-specific neurons that express *Vsx1* include both Notch^ON^ and Notch^OFF^ fates^27^. Among domain-ignoring neurons, both the Notch^ON^ Mi1 fate and the Notch^OFF^ T1 fate lack Vsx1, while domain-independent Vsx1 expression occurs in Notch^OFF^ types such as Dm2 and Tm5c and in Notch^ON^ types such as Tm24 and Tm25, in different temporal contexts (**Extended Data Figure 4A**)^25,27,46,47^. Consistently, Coyne and colleagues showed that the *Dm2-2* fragment contains subregions with accessibility specific to distinct temporal and Notch identity combinations, suggesting that element-internal regulatory logic, rather than a specific Notch signaling status, determines which fates express *Vsx1*^29^.

Together, these results indicate that engagement with the spatial re-expression program is a property of a neuron’s fate identity, specified by the intersection of temporal and Notch inputs. Because spatial identity in the medulla is carried at a single locus rather than as a genome-wide program, cells from different domains already share nearly identical molecular states. Domain independence therefore does not require reconciliation of two distinct programs but only modulation of one locus. For fates that exclude *Vsx1*, this includes loss of accessibility at the bookmarked *Vsx1/2 locus*, as observed in Mi1 and Tm1/2/4/6. For fates that express *Vsx1* domain-independently, *Vsx1* is incorporated into the temporal-Notch program through fate-specific regulatory elements like any other terminal selectors, so that its expression no longer depends on the domain-bookmarked program in the *Vsx1/2* locus but instead on specific enhancer elements like the *Dm2-2* element. In both cases, domain-ignoring fates are only specified by temporal and Notch inputs, without requiring the domain-restricted chromatin trace, a regulatory logic reminiscent of Optix-domain fates.

## Discussion

### *Drosophila* medulla as a model for spatial memory without relay

Our results reveal that spatial identity in the medulla is encoded in persistent chromatin accessibility only at the locus of the spatial factors themselves. Progenies from the Vsx and Bi domains retain a minimal trace that is concentrated at the same genes whose re-expression in postmitotic neurons executes domain-specific fates. PRC2-mediated silencing maintains domain restriction of Vsx1 re-expression, while Notch signaling and temporal identity jointly determine which neurons engage or bypass the spatial program. This strategy is in contrast to the *Drosophila* ventral nerve cord, where genome-wide chromatin remodeling in progenitors encodes spatial identity^10,11^, or the vertebrate neural tube, where spatial information is relayed via downstream effectors before the original factors disappear^14^. While chromatin accessibility differences among ventral spatial domains of the neural tube are also less prominent than that among temporal identities^48,49^, unlike the medulla, spatial regulation in the neural tube still involves thousands of differentially accessible regions rather than being concentrated at a single spatial-factor locus, underscoring the unusually restricted scope of the medulla’s spatial trace.

This PRC2 dependency in the *Drosophila* medulla is selective: among the factors we tested, only *Vsx1* was derepressed upon *E(z)* loss, consistent with *Vsx1* being responsive to a broadly present neuronal activator that PRC2 ordinarily restrains. Because the cross-repressive network that initially excludes *Vsx1* from non-Vsx neuroepithelium dissolves at the progenitor transition, when spatial-factor proteins are no longer detectable^17^, PRC2-mediated silencing maintains repression in the absence of this initiating input. *Bi*, despite a similar expression trajectory and H3K27me3 occupancy, is not derepressed upon *E(z)* loss, consistent with its known dependence on spatially restricted Wg signaling^19,50^, which may preclude de-repression by PRC2 removal alone. Although our transcriptomic and chromatin analyses did not detect domain-enriched transcriptional regulators in progenitors that could relay spatial information, post-transcriptional or protein-level mechanisms not captured by these assays cannot be excluded.

What maintains chromatin accessibility at the *Vsx1/2* and *Bi* loci in their home domains through progenitor stages? The specific differentially accessible elements at these loci largely differ across developmental stages, indicating that what persists is not a fixed set of regulatory elements but a locus-level chromatin state. Ongoing weak transcription at both *Vsx1/2* and *Bi* nuclear loci may establish and maintain a chromatin state by resisting PRC2 silencing^43,44,51,52^. Prior neuroepithelial expression alone does not suffice: *Optix* is transcribed in the neuroepithelium but shows no persistent domain-specific accessibility and is not re-expressed in neurons. Both *Vsx1/2* and *Bi* share this locus-level architecture, consistent with ongoing transcription at spatial-factor loci producing a permissive chromatin state in home-domain progenies. Why sustained transcription at these loci fails to yield detectable product in progenitors remains unresolved and would require direct measurement of transcript fate.

While re-expression of *Vsx1/2* or *Bi* confers domain-specific fates, *Optix* itself is not expressed in any medulla neuron and we have not identified a single terminal identity program shared by Optix-domain progenies^27^. At the level of the *Vsx1/2* and *Bi* spatial programs examined here, however, multiple perturbations convergently generate Optix-domain or Optix-like fates: loss of *Vsx1/2* shifts neurons toward Optix-domain fates, and loss of *Bi* shifts TE neurons toward the domain-ignoring Mi1 fate. In contrast ectopic expression of either *Vsx1* or *Bi* promotes domain-specific identities at the expense of Optix-domain or Optix-like fates. This suggests that Optix-domain fates are specified by temporal and Notch programs, resulting in a ground-state output from which spatial factor re-expression diverts neurons into domain-specific alternatives.

Domain-ignoring fates similarly achieve independence from spatial origin either by excluding spatial-factor re-expression or by recruiting it through fate-specific regulatory elements. Optix contributes to this architecture at the neuroepithelial stage, where mutual cross-repression among spatial factors establishes domain boundaries^16,38^: *Optix* represses *Vsx1* expression outside the Vsx territory, setting the initial transcriptional state at spatial-factor loci that the single-locus chromatin trace then carries through progenitor stages.

### Columnar stoichiometry creates a numerical constraint on spatial patterning

Precise but minimal encoding of spatial identity makes overriding spatial patterning structurally inexpensive, but why do specific neuron types exercise this option? Spatial subdivision creates a numerical constraint: a fate restricted to one progenitor domain is generated from a smaller progenitor pool, whereas repetitive columnar organization in the visual system requires some visual neuron types to maintain 1:1 stoichiometry with individual ommatidial inputs across the medulla. All six major first-relay targets of L1 and L2, the lamina ON and OFF input channels for elementary motion detection, are domain-ignoring: Mi1 and Tm3 are connected to L1, and Tm1, Tm2, Tm4, and T1 to L2^53,54^. Producing these types across multiple spatial domains expands the progenitor pool available to generate them and provides one route to achieving their 1:1 columnar stoichiometry, while other neurons achieve comparable stoichiometry despite remaining predominantly domain-specific^27,53,54^, presumably meeting the same numerical demand through alternative developmental strategies such as production during extended temporal windows or reduced apoptosis. Thus, spatial bypass is one developmental solution to the abundance constraint imposed by spatial subdivision, but not the only one.

Spatial bypass is not confined to these motion-channel relays. Dm2 and Tm5c, associated with luminance and color processing, likewise adopt equivalent identities across spatial domains^53,55,56^, but their ∼1:1 columnar stoichiometry^54^ uses a different strategy to reach the same quantitative constraint: Unlike the L1/L2 targets that avoid domain-specific *Vsx1/2* re-expression, these types instead express *Vsx1* through fate-specific regulatory programs independently of spatial origin.

The occurrence of bypass in different circuits and through distinct regulatory mechanisms therefore supports a broader architectural conclusion: single-locus bookmarking makes spatial restriction optional for individual fates.

### Minimal encoding enables intrinsic flexibility

The antero-posterior axis of the optic lobe examined in this work is one of several spatial axes patterning the medulla. The dorso-ventral axis is defined by neuroepithelial expression of ventrally expressed *Disco* and dorsally expressed *Spalt*^57^, which, like *Optix*, are only transiently expressed but are not re-expressed in a domain-specific fashion in neurons. Furthermore, the Optix domain is further subdivided into smaller domains by TFs like *Brk*^15,30^. How the specific spatial information from dorso-ventral patterning and Optix subdivision is maintained and translated into distinct fates remains elusive. A related question is emerging in the vertebrate neural tube, where one recent lineage analysis suggests that the canonical dorsoventral progenitor domains are nested within a smaller number of broader lineage compartments^58^. Together with evidence that spatial determinants are interpreted through a dynamically regulated chromatin landscape^59^, these findings emphasize the need to understand how individual spatial distinctions are encoded and how strongly that encoding constrains neuronal fate.

Taking all axes into consideration, most medulla neuron types ignore at least one spatial axis. For example, the motion pathway neurons described above ignore the Vsx-Optix boundary, the dorso-ventral axis, and intra-Optix boundaries^27^, while Mi4 and Mi9 only respect the Vsx-Optix boundary but are produced uniformly along the dorso-ventral axis and across intra-Optix boundaries. If boundaries along all spatial axes were universally respected, the combinatorial intersection of these axes would have the capacity to subdivide the medulla into far too many fates. That most types selectively engage only a subset of available axes suggests that the developmental architecture allows individual neuron types to opt in or out of spatial patterning as needed.

The minimal scope of the chromatin trace for the Vsx and Bi domains may be central to allowing individual neuronal fates to bypass spatial restrictions. The tradeoff between spatial diversification and the need for specific neuron types to escape domain boundaries is not unique to the medulla: neurons of the medial motor column of the vertebrate spinal cord are generated across *Hox*-defined segments throughout the spinal cord^60^. However, even this closest parallel is partial: while the medulla produces dozens of fully domain-ignoring types, medial motor column neurons retain segment-specific *Hox* expression and likely serve partially distinct roles. How readily a developing nervous system can accommodate such exceptions may depend on how deeply spatial identity is encoded: systems that defer execution to minimal, late-acting cues create room for structured flexibility, while those that lock spatial programs into elaborate transcriptional or chromatin cascades may find it harder to make exceptions. In the medulla, single-locus chromatin memory therefore preserves spatial information while allowing each neuronal fate to engage it only when needed.

## Acknowledgements

We thank Y.-C. D. Chen, N. Konstantinides, J. A. Malin, and R. Rajesh for thorough feedback on the manuscript and members of the Desplan lab for helpful discussions; R. Loker for his insights on Polycomb-mediated regulation and pointing us to relevant datasets; T. Erclik for insights into Vsx1 regulation; G. Struhl for advice on Bi perturbation; M. Sato for sharing a Runt driver line; the Bloomington Drosophila Stock Center for fly stocks (NIH P40OD018537); FlyBase for comprehensive curation of genetic and reagent information (NHGRI U24HG013300); and the Genomics Core at the NYU Center for Genomics and Systems Biology (GenCore) for scientific support and sequencing services. Computational work was supported in part by the resources and personnel of the NYU High-Performance Computing Core and by Google Cloud through the Research Credits Program (award GCP19980904).

## Author contributions

Y.-C.C. and C.D. conceived the project. Y.-C.C. performed experiments and analyses. S.N. contributed to Notch loss-of-function experiments and characterization of domain-ignoring types. B.J.C. designed the somatic CRISPR mutagenesis system and generated the CRISPR lines and low-perdurance GFP reporter used in reporter assays. A.B. validated somatic CRISPR for Bi. M.N.Ö. identified Vsx-Fragment 2. M.N.Ö. and R.C. contributed to the interpretation of Dm2-2 regulation. C.D. supervised the project. Y.-C.C. and C.D. wrote the manuscript with input from all authors.

## Funding

The Desplan laboratory was supported by grants from the National Eye Institute R01EY017916 and R01EY013010 to C.D. Y.-C.C. was supported by New York University through the MacCracken Fellowship, by an NYSTEM institutional training grant (Contract C322560GG), and by a Scholarship to Study Abroad from the Ministry of Education, Taiwan. S.L.N. was supported by a DURF fellowship from NYU. B.J.C. was supported by NIH fellowship F32EY027682. M.N.Ö. and R.C. are supported by the National Institute of Neurological Disorders and Stroke (NIH R00NS125117 to M.N.Ö.) and Stowers Institute for Medical Research. A.B. was supported by Tamkeen under the NYU Abu Dhabi Research Institute Award to the NYUAD Center for Genomics and Systems Biology (ADHPG-CGSB).

## Methods

### Drosophila melanogaster strains and husbandry

See Supplementary Table 4 for the genotype used for each figure and a complete list of stocks used. Flies were grown on standard cornmeal, molasses, and yeast media at 25°C on a 12-hour light/dark cycle unless otherwise specified. Both male and female animals were analyzed for each experiment, but no sex-specific differences were noted. Late third-larval instar larvae were defined as larvae wandering on the vial wall.

### Generation of transgenic animals

Transgenic constructs were cloned into the following plasmids: *Act5C-FRT-STOP-FRT-Cas9-2A-myrTomato* (pAct-FSF-Cas9_t2aTom; *VK05* (BDSC 9725)), *UAS-nls::sfGFP::PEST-T2A-B2ase* (pUASsfGFPPEST_t2a_B2ase; *attP1* (BDSC 34760)), *U6-Bi sgRNA* (pCFD5-white; *attP2* (BDSC 8622)), *UAS-Vsx1[CDS]::FLAG* (pJFRC177-10XUAS-SapI; *ZH-86Fb* (BDSC 24749)), *UAS-Bi[CDS]::FLAG* (pJFRC177-10XUAS-SapI; *ZH-86Fb*), *Vsx-Frag-1-Gal4* (pBPGUw-MCS; *attP2*), *Vsx-Frag2-Gal4&lexA* (pBPGUw-bidirectional; *attP2*) and *Vsx-Frag-3-Gal4* (pBPGUw-MCS; *attP2*). See Appendix 1 for plasmid maps. All constructs except *Vsx-Frag2-Gal4&lexA* were injected and integrated into corresponding attP sites by BestGene (Chino Hills, CA, U.S.A.), while *Vsx-Frag2-Gal4&lexA* was injected and integrated by WellGenetics (Taipei, Taiwan).

For guide RNAs, we used the design strategy reported by Choi *et al.*^61^. Briefly, six sgRNAs were driven by the ubiquitous U6:3 promoter using the pCFD5 multiplexing strategy. These sgRNAs targeted three sites in the first coding exon, with each target site represented twice in the sgRNA array (**Extended Data Figure 2H**). The coordinates for the *Vsx* fragments are: Fragment 1: chrX:5585828-5587942; Fragment 2: chrX:5582776-5584000; Fragment 3: chrX:5576167-5579314.

### Temperature-sensitive neuronal perturbations

For neuronal gain- and loss-of-function experiments, transgenes were expressed using *elav^C155^-Gal4* with a temperature-sensitive Gal80 repressor (*tub-Gal80ts*). Crosses were maintained at 18°C, and larvae were shifted to 29°C for the final 48 hours of larval development before dissection at the wandering third-instar stage. At 29°C, this shift approximately spans the third larval instar, when medulla neurons are generated from the outer proliferation center. For loss-of-function experiments, *UAS-mCherry RNAi* was used as a control to account for engagement of the endogenous RNAi machinery. For gain-of-function experiments, *UAS-LacZ* was used as a control to account for the effects of transgene expression on proteostasis. See Supplementary Table 4 for complete genotypes.

### MARCM clonal analysis

Mosaic analysis with a repressible cell marker (MARCM) was used to generate *Su(H)^del47^* mutant clones in the medulla. Larvae carrying hs-FLP, MARCM components, and the *Su(H)^del47^ FRT40A* chromosome (or *FRT40A* alone for controls) were heat-shocked in a 37°C water bath for 30 minutes, three days before dissection at the wandering third-instar stage, corresponding to late second instar to early third instar. Clones were identified by GFP expression. Vsx-domain clones were identified by their position in the anterior medulla and confirmed by the presence of flanking non-clonal cells expressing Vsx2. Only Hth-positive neurons within clones were scored. Vsx2-positive Hth-positive neurons were counted as a proportion of all Hth-positive neurons within Vsx-domain clones. Clones from the same animal were pooled before proportion calculation, so that each animal constituted one biological replicate. Vsx2 was used as a proxy for *Vsx1/2* expression because Vsx1 and Hth antibodies were raised in the same host species and could not be used simultaneously; Vsx1 and Vsx2 are co-expressed in Vsx-domain neurons at the third larval instar (**Extended Data Figure 4B**).

### Somatic CRISPR for *Bi* loss of function

Somatic disruption of *Bi* was achieved using a FLP-inducible Cas9 cassette (*Act5C-FRT-STOP-FRT-Cas9-2A-myrTomato*) combined with a U6-driven tandem sgRNA construct targeting the first coding exon of *Bi* (**Extended Data Figure 2H-I**). Larvae were heat-shocked in a 37°C water bath for 1 hour, three days before dissection at the wandering third-instar stage, corresponding to late second instar to early third instar. Heat-shock-induced FLP excised the transcriptional stop cassette, enabling Cas9 and myrTomato co-expression in clones. Clones were identified by myrTomato fluorescence. Because Cas9 expression was driven by the ubiquitous *Act5C* promoter beginning soon after clonal induction, *Bi* disruption was not restricted to neurons and could affect neuroepithelial cells, neuroblasts, ganglion mother cells, and neurons.

Control animals carried the Cas9 cassette without sgRNAs. TE neurons (*Dimm^+^ Bsh^+^*) were scored as a proportion of all *Bsh^+^* cells within myrTomato-positive clones, restricted to the TE territory defined by kernel density estimation on control samples (see Region of interest definition). Two independent RNAi lines targeting *Bi* were tested and failed to achieve knockdown in optic lobe neurons (**Extended Data Figure 2E-G**), motivating the somatic CRISPR approach.

### Immunohistochemistry

Fly optic lobes were dissected in chilled Schneider’s medium (ThermoFisher, 21720024). Dissected samples were transferred to 4% paraformaldehyde in Schneider’s medium and fixed at 4°C for 15 minutes. After 3 washes in 1% PBTx (1% TritonX-100 (ThermoFisher, A16046-AE) in PBS (Growcells, MRGF6395)) for 15 minutes, samples were blocked in blocking buffer (5% horse serum (Gibco, 26050070) in 1% PBTx) for at least 1 hour at room temperature. See Supplementary Table 5 for primary antibodies and titers used in this study. Primary antibody mix was prepared by diluting each antibody at its specified titer in blocking buffer and was added to the sample to incubate on a horizontal shaker at 4°C for two nights. The samples were then washed 3 times for 15 minutes in 1% PBTx at room temperature and incubated in secondary antibody mix on a horizontal shaker at 4°C for two nights. Finally, the brains were washed 3 times for 15 minutes in 1% PBTx at room temperature, rinsed in PBS twice, and then mounted in Slowfade (ThermoFisher, S36396) before imaging with a Leica SP8 or Stellaris confocal microscope using a 63x glycerol objective.

### Hybridization Chain Reaction Fluorescence *In Situ* Hybridization (HCRv3)

Fly optic lobes were dissected in chilled Schneider’s medium. Dissected samples were transferred to 4% paraformaldehyde in Schneider’s medium and fixed at room temperature for 20 minutes. After 3 washes in 1% PBTx, samples were equilibrated in Probe Hybridization Buffer (Molecular Instruments) for 10 minutes at 37°C. See Appendix 2 for probe sets used in this study. Customized probe sets were designed with Insitu Probe Generator (RRID:SCR_025981)^62^.

1 µM probe was diluted 1:250 in Probe Hybridization Buffer to prepare the probe mix, and samples were incubated in probe mix at 37°C overnight. Samples were then washed with Probe Wash Buffer (Molecular Instruments) for 10 minutes at 37°C 4 times and twice in 5x SSCT (5xSSC (Growcells, 75235) + 0.1% v/v Tween-20 (Biorad, 1610781)) for 5 minutes. Samples were then equilibrated in Amplifier Buffer for 10 minutes at room temperature. Amplifiers (Molecular Instruments) were denatured by a thermocycler to 95°C for 90 seconds and allowed to cool for 30 minutes at room temperature, protected from light. Amplifier mix was prepared by diluting each Amplifier hairpin 1:50 in Amplifier Buffer. Samples were incubated in amplifier mix at room temperature overnight protected from light. Samples were then washed with 5x SSCT 3 times for 10 minutes at room temperature and optionally incubated in Hoechst33342 (ThermoFisher, 62249) diluted 1:500 in 5x SSCT at room temperature overnight. Samples were then rinsed in PBS twice and mounted in Slowfade before imaging with a Leica SP8 or Stellaris confocal microscope using a 63x glycerol objective.

#### RFP signal boosting in *Bi* loss-of-function HCR experiments

In *Bi* loss-of-function HCR experiments, RFP signal was boosted by adding an Alexa Fluor 568-conjugated nanobody (ProteinTech, rb2AF568) during the Hoechst33342 incubation at a 1:500 dilution.

### Image segmentation and mask generation

Image stacks were processed using Cellpose v4.0.9^63^ to generate three-dimensional segmentation masks. TIFF images were analyzed with the Cellpose-SAM model, with the channel axis set to 1 and the z axis set to 0. The xy pixel size was extracted from Fiji metadata, and the z-step was read from the ImageJ TIFF spacing metadata. These values were used to calculate image anisotropy as z-step divided by xy pixel size before segmentation.

Cellpose was run in 3D mode with a batch size of 64, minimum object size of 100, cell probability threshold of 0, flow threshold of 0, stitch threshold of 0.25, flow smoothing of 0, and tile normalization disabled. The resulting labeled masks were then post-processed using region-property measurements. Objects were removed if their asphericity exceeded 2, their mean intensity in the marker channel used for segmentation was below the 2nd percentile of detected objects, or their convex area was less than 500 pixels/voxels. Labels failing these criteria were set to background, producing the final filtered mask used for downstream analysis.

Filtered masks were then manually curated via Cellpose GUI to remove non-optic lobe cells and add missing cells when necessary. Curated cell masks were then treated as labeled 3D ROIs and measured against the corresponding signal image. To analyze cells in a shared standardized coordinate system, we relied on the fact that the larval medulla cortex is a shallow dome-shaped structure, and principal component analysis therefore tends to capture the sagittal plane with the first and second components. For each sample, cell centroids were projected onto three principal-component axes. The first two PCA axes were then manually rotated by a sample-specific angle so that the positive direction of the x-axis is anterior. The rotated coordinates were then scaled to a common [-1, 1] range, producing the aligned x_std and y_std coordinates used for spatial comparisons.

Marker-positive cells were assigned using experiment-specific intensity thresholds manually set from the distribution of negative/background cells and held fixed across control and perturbation samples within each experiment. No explicit depth-dependent intensity correction was applied; marker thresholds were therefore set within each experiment using samples imaged under identical settings.

#### Region of interest definition for TE neurons in *Bi* somatic mutant clones

The TE-region ROI was defined from Cas9-only control cells that were double-positive for *Dimm* and *Bsh*. A two-dimensional kernel density over the aligned PCA coordinates was estimated with bandwidth selected using Silverman’s rule. A 50% maximum kernel density cutoff was used to create a binary mask. Cells were retained for the TE-region analysis when their aligned coordinates fell within the mask.

#### Radial position analysis for Vvl and Toy

For Vvl and Toy, radial position was calculated from the aligned coordinates as the distance from the center of the normalized PCA plane. This reduced each cell’s position to a single radial coordinate while preserving whether cells were concentrated near the center or spread toward the periphery. For each biological sample, the variance of radial position was calculated, and the per-sample variances were compared between *mCherry* RNAi and *E(z)* RNAi conditions with a two-sided Welch’s t-test.

#### Vsx1 expression ratio within GFP reporter expressing cells

For the *Vsx* reporter analysis, GFP+ cells were segmented and assigned to a spatial ROI using their angular position in the aligned coordinate system. Within this ROI, each reporter-positive cell was classified as Vsx1-positive or Vsx1-negative. As the reporter experiments were done separately and imaged with different laser settings, intensity cutoff for the Vsx1 channel was manually determined per reporter by inspection of the intensity histogram and anatomical information. The Vsx1 expression ratio was then summarized per biological sample as the number of Vsx1-positive reporter cells divided by the total number of reporter-positive cells in the ROI. The conservative Vsx-domain ROI was defined as angular coordinates from −π/10 to π/10 in the aligned coordinate system.

### Quasibinomial test for cell counts

For proportion-based comparisons, cells were first summarized per biological sample as a numerator and denominator, such as marker-positive cells over total eligible cells. Conditions were compared with a quasibinomial GLM using the sample-level proportion as the response and the denominator as the weight (using glm() in R v4.5.1 with the quasibinomial family). The quasibinomial model was used instead of a simple binomial test because cell counts within a sample are not fully independent and biological samples can show extra-binomial variation; the model estimates an overdispersion parameter while retaining denominator information.

For each perturbation, cell fates were scored as proportions within a temporal-cohort denominator, so that changes reflect fate shifts within a defined progenitor pool rather than global cell number changes. Spatial restrictions were applied when marker combinations were ambiguous outside the domain of interest.

For *Vsx1/2* perturbations in the Run+ cohort, Pm4 identity (Ct**-**Run**+**) was scored as a proportion of all Run^+^ cells restricted to the anterior half of the medulla cortex (standardized x > 0). Posterior Run^+^ Ct**^-^** cells were excluded because this marker combination does not uniquely identify Pm4 outside the Vsx domain. For *Vsx1/2* perturbations in the *Ey^+^ Kn^+^* cohort, Tm5e identity (*Ey^+^ Kn^+^ D**^-^***) was scored as a proportion of all *Ey^+^ Kn^+^* cells restricted to the anterior half of the medulla cortex (standardized x > 0).

For *Bi* loss of function, TE identity (*Dimm^+^ Bsh^+^*) was scored as a proportion of all *Bsh+* cells within myrTomato-positive clones, restricted to the TE territory defined by kernel density estimation on control samples (see Region of interest definition). For *Bi* gain of function, TE (*Dimm^+^ Bsh^+^*) was scored as a proportion of all Bsh^+^ cells in the anterior half of the medulla cortex (standardized x > 0), where TE neurons are not detected normally.

For *E(z)* knockdown, Vsx1 expansion was scored as the fraction of all Vsx1^+^ neurons located in the posterior half of the medulla cortex (standardized x < 0), representing Optix- and Bi-domain progenies. Bi expansion was scored as the fraction of all Bi^+^ neurons located in the anterior half (standardized x > 0). For Toy and Vvl, the radial variance of marker-positive neurons, which reflects the thickness of the expression domain, was compared between conditions (see Radial position analysis).

For *Su(H)* MARCM, Vsx2^+^ Hth^+^ neurons were scored as a proportion of all Hth^+^ neurons within Vsx-domain clones. Clones from the same animal were pooled before proportion calculation; each animal constituted one biological replicate.

### Domain-resolved single-cell multiome sequencing

To simultaneously profile Vsx-, Optix-, and Bi-domain progenies, we crossed nuclear GFP reporters driven by *pxb^MI05058^-T2A-Gal4* (Vsx), *Optix^NP2631^-Gal4* (Optix), and *dpp-Gal4* (Bi) (**Figure 2A**) with a selected set of isogenic and whole-genome sequenced strains in the *Drosophila* Genetic Reference Panel (DGRP lines: 40, 208, 235, 307, 313, 315, 320, 382, 383, 391, 399, 406, 427, 441, 492, 505, 517, 852, 897). For each experiment, males from each reporter line were mated with virgins collected from a unique set of DGRP lines, so cells from the F1 offspring of each reporter can be distinguished by line-specific single nucleotide polymorphisms^26^.

For sample collection, we reared the animals at 25°C and dissected both male and female wandering larvae to retrieve larval optic lobes by removing the ventral nerve cord and the central brain. Dissection was done for at most one hour. Optic lobes were kept in ice-cold Schneider’s medium until dissociation. Optic lobes were moved to dissociation buffer (2 mg/mL Dispase (Sigma, D4693-1G) + 2 mg/mL collagenase (Sigma, C0130) in Schneider’ s medium) and incubated at 26°C for 30 minutes. Samples were then gently rinsed with PBS + 1%BSA (126615, Millipore) twice. In 100 µL 1% BSA, samples were mechanically dissociated by pipetting vigorously 50 times with a P200 pipet on ice without introducing bubbles. Cell suspension was then sieved through a pre-wetted 10µm pluriStrainer (pluriSelect, 43-10010-40) into a LoBind mL tube (Eppendorf, 022431021) rinsed with 1% BSA on ice. The tube for dissociation was then washed twice with 50µL 1% BSA, and the wash buffer was sieved through pluriStrainer to recover as many cells as possible.

The cells were then sorted by FACS for GFP-positive cells into another LoBind 1.5mL tube rinsed with 1% BSA and containing 100µL buffer in the tube, with gating set via negative control (Canton-S optic lobe dissociated with the same protocol above). Sorted cells were spun down at 800 x g with a swing-bucket centrifuge (Eppendorf 5810R) at 4°C. Supernatant was removed with 10 µL left, and cells were resuspended with 90 µL lysis buffer added (10mM Tris-HCl (Sigma T2194), 146mM NaCl, 1mM CaCl_2_, 21mM MgCl_2_ (ThermoFisher, AM9010), 0.03% Tween-20 (Biorad, 1662404), 1mM DTT (ThermoFisher, P2325), 1U/mL Protector RNase Inhibitor (Sigma, 3335402001), 0.01% BSA (126615, Millipore), 5% Ez Lysis Buffer (Sigma, NUC101-1KT)). The suspension was gently pipetted with a P200 pipette with a wide-bore tip 10 times and incubated on ice for 10 minutes. 400 µL of wash buffer (10mM Tris-HCl (Sigma T2194), 10mM NaCl, 3mM MgCl_2_ (ThermoFisher, AM9010), 1mM DTT (ThermoFisher, P2325), 1U/mL Protector RNase Inhibitor (Sigma, 3335402001), 0.01% BSA (126615, Millipore)) was added, and cell suspension was centrifuged at 800 x g at 4°C for 5 minutes. Then, 490 µL supernatant was removed, and 100 µL permeabilization buffer (wash buffer + 0.01% digitonin (ThermoFisher, BN2006)) was added carefully without resuspension. The sample was then incubated on ice for 5 minutes. Finally, the sample was centrifuged at 800 x g at 4°C for 5 minutes, and as much supernatant as possible was removed without disrupting the pellet, which might not be visible. The pellet was resuspended in 7 µL diluted nuclei buffer (10x Genomics).

We then used Chromium Next GEM Single Cell Multiome ATAC + Gene Expression kits (10x Genomics, PN-1000283) following the manufacturer’s instructions. Briefly, the nuclei were tagmented via Tn5 transposition at 37°C for 1 hour and loaded into a Chip J (10x Genomics, PN-1000230) for generation of gel bead-in-emulsions (GEMs) using a Chromium Controller. The GEMs were cleaned up, pre-amplified for 7 cycles and split into ATAC and RNA samples. ATAC libraries were constructed with 9 cycles of sample index PCR, while RNA libraries were constructed via 9 cycles of cDNA amplification, fragmentation and end-repair of 20µL amplified cDNA, adapter ligation, and 15 cycles of sample index PCR. Four libraries were generated from 3 independent experiments (the last experiment generated two libraries), with each library aiming for the recovery of 15,000 cells. The libraries were subjected to paired-end Illumina sequencing using NovaSeq 6000: 50x8x24x49 cycles for ATAC libraries and 28x10x10x90 cycles for RNA libraries.

### Whole-genome sequencing for reporter genotyping

F1 hybrids were collected from crosses between males from each reporter line and virgins collected from the dm6 reference genome line (BDSC 2057). Genomic DNA was extracted with Quick-DNA Tissue/Insect Miniprep Kit (Zymo Research, D6016) and constructed into libraries with NEBNext UltraII FS DNA Library Prep Kit (New England Biolabs, E7805S) and sequenced with NovaSeq 6000 at 100M reads per library. Reads were adapter-trimmed with cutadapt and aligned to the Ensembl release 88 dm6 genome with bwa-mem2. Alignments were filtered at mapping quality 30, sorted, indexed with samtools, and processed with Picard to add read groups and mark duplicates. Variants per reporter line were called with two complementary workflows:

1. GATK HaplotypeCaller was run per chromosome, followed by joint genotyping, single nucleotide variant (SNV) and indel hard filtering.
2. bcftools mpileup/call was used to call variants from the same BAMs.

Variants passing the bcftools quality filter were intersected with the GATK calls to define a conservative reporter-line variant set. The resulting reporter-line VCF was used to remove driver-confounded SNVs from the genetic barcoding reference.

### *In silico* Demultiplexing of multiome data

Genetic barcodes were assigned from single nucleotide variants (SNVs) that distinguished the DGRP lines used in each multiplexed library. Briefly, DGRP variants were first restricted to the selected parental lines, filtered to informative SNPs that are only present in one of the DGRP lines used in an experiment and are homozygous, and compared with reporter-line variants so that driver-confounded sites were excluded. For each library, expected F1 genotypes were encoded from the DGRP parent and its spatial domain of origin^26^.

To reduce computational load, cell barcodes were permissively filtered to remove those with fewer than 200 detected genes or fewer than 300 ATAC fragments. The top 100 expressed genes in each library were masked from the variant set to reduce RNA-expression bias, and variants were further restricted to sites with at least 10 supporting reads in the RNA or ATAC BAMs.

RNA and ATAC alignments were merged for each library, pileups were generated with dsc-pileup, and cell identities were assigned with demuxlet using the genotype field^31^. freemuxlet was also run as a genotype-free check. Downstream analyses retained cells assigned as singlets by demuxlet or freemuxlet, provided that the inferred DGRP-to-spatial-origin label mapping was consistent between methods.

### Analysis of domain-resolved single-cell multiome data

#### Preprocessing with cellranger-arc

Single-cell multiome libraries were processed with cellranger-arc v2.0.2 using a custom Ensembl release 88 dm6 reference with Stinger added to the reference genome as an extra contig. For each library, cellranger-arc count produced gene expression matrices, ATAC peak matrices, aligned RNA and ATAC BAMs, fragment files, and initial peak calls. The RNA matrix was extracted from the raw multiome output and processed with CellBender remove-background (v0.3.2)^64^; the CellBender-corrected gene expression matrix was used for downstream Seurat object construction.

#### Metadata curation

RNA counts, ATAC insertion counts, demultiplexing calls, library labels, DGRP line identities, and spatial origin labels were assembled into a Seurat object with a Signac chromatin assay^65,66^. Gene features were restricted to annotated genes on standard chromosomes, ribosomal RNA genes were removed, and ATAC peaks overlapping the modEncode dm6 blacklist or non-standard chromosomes were excluded. Cell names were prefixed by library to keep barcodes unique across libraries.

#### Quality control and data filtering

Cells were retained if they passed the singlet consistency filter (see *In silico* Demultiplexing of multiome data), had more than 200 RNA UMIs and more than 300 ATAC insertion events, and were confirmed by manual marker review not to be low-confidence or non-optic-lobe clusters. ATAC-specific quality control was then performed with Signac, retaining cells with TSS enrichment greater than 2 and nucleosome signal less than 4.

#### Consensus peak definition

Consensus ATAC peaks used for the final chromatin assay were defined from cluster-level insertion profiles rather than from per-library Cell Ranger ARC peak calls. After manual removal of low-confidence clusters, cells were grouped by retained WNN cluster. For each cluster, ATAC fragments from all libraries were pooled and converted to Tn5 insertion sites. MACS2 callpeak was run on each cluster-level insertion BED with --nomodel, --shift -75, --extsize 150, --keep-dup all, --call-summits, and -q 0.05 using the dm6 genome size.

Cluster summit calls were concatenated and sorted by MACS2 summit score. A greedy consensus set was then selected by retaining higher-scoring summits first and discarding lower-scoring summits less than 150 bp from an already retained summit on the same chromosome. Retained summits were expanded symmetrically by 150 bp to produce 300-bp intervals; summits whose expanded intervals would cross chromosome boundaries were excluded. The resulting intervals were sorted by genomic coordinate and used to quantify ATAC insertions for downstream analyses. Non-standard chromosomes and intervals overlapping the dm6 blacklist were removed during Seurat object construction.

#### Dimensionality reduction and clustering

RNA counts were normalized with SCTransform while regressing mitochondrial percentage, and ATAC counts were normalized with TF-IDF followed by latent semantic indexing. Variable genes and peaks were selected iteratively to preserve subtly diversified populations and differences between spatial origins. Briefly, global variable features were selected and used for dimensionality reduction and clustering. The variable feature list was then regenerated by appending: (1) differentially enriched features among spatial origins per cluster, and (2) variable features found per cluster. This process was repeated for five iterations and features present in three or more iterations were retained for the final analysis.

After filtering, RNA PCA and ATAC LSI embeddings were corrected for batch effects (library of origin) with Harmony^67^, and a weighted nearest-neighbor graph was built from both modalities using 120 PCs and 110 LSI dimensions with 20 nearest neighbors. Dimension choices were guided by neighborhood-stability analyses, and graph clustering was evaluated across resolutions using eigengap and silhouette diagnostics. The final clusters were visualized with UMAP.

#### Cluster annotation

Clusters were annotated from aggregate marker-gene expression and reference-guided preliminary labels assigned by a published neural network classifier trained on gene expression data^25^. Major cell classes were assigned using markers for neuroepithelial cells, neuroblasts, ganglion mother cells, neurons, lamina precursor cells, and glia. Temporal identities for progenitors were assigned from temporal transcription-factor expression, grouping cells into early- or late-born cohorts, while temporal identities of neurons were assigned from known postmitotic regulators^20,22–24^, and neuronal Notch status was annotated via the expression of Ap^20^.

#### Visualization of ATAC-seq

Coverage tracks were generated by splitting ATAC BAMs by annotated groups with scbamop (https://github.com/chenyenchung/scbamop), converting the split BAMs to CPM-normalized bigWig files with deepTools^68^, and averaging tracks across libraries. bigWig files were then visualized with pyGenomeTracks^69^. For Figure 5C, four pseudobulk neuronal groups were defined from Vsx-domain neurons in the multiome dataset. The Vsx-domain neuron track included all Vsx-origin neurons except those annotated as Hth/Notch^ON^ (Mi1) or Hth-Opa/Notch^ON^ (Tm1/2/4/6). The Mi1 and Tm1/2/4/6 track comprised Hth/Notch^ON^ and Hth-Opa/Notch^ON^ neurons from the Vsx domain. The D/BarH1 Notch^OFF^ and Notch^ON^ tracks comprised Vsx-domain neurons of the corresponding temporal-Notch identity; the Notch^OFF^ cohort includes Dm2 among multiple other types.

### Identity Segregation Score

Identity Segregation Score (ISS) was used to quantify whether cells with the same biological identity were locally enriched in the multimodal neighborhood graph. Simpson concentration was estimated using the unbiased distinct-pair estimator. For identity counts (*n_i_*) among (*N*) cells, the global Simpson index was^70,71^:

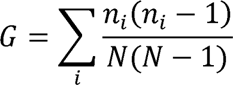

For each cell (*c*), let *m*_*ci*_ denote the number of neighbors with identity *i*, and let Σ_*c*_ = Σ_*i*_*m_ci_* be the number of its 20 nearest neighbors with non-missing identity labels. The local Simpson index was:

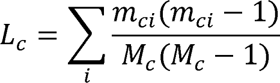

Neighborhoods containing fewer than two non-missing labels were assigned a missing score. ISS was then calculated as:

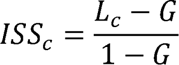

Under random mixing, the expected ISS is zero. Positive values indicate local segregation of the identity, while negative values indicate greater local mixing than expected from the global identity composition. Spatial ISS was calculated across the complete L3 multimodal neighborhood graph using spatial-origin labels and summarized separately by cell class. For temporal ISS, the global reference population comprised NBs, GMCs, and neurons with temporal annotations; scores were subsequently analyzed separately for these classes.

Spatial and temporal ISS values were compared within each stage using paired per-cell differences, (*D* = *ISS*_*spatial*_ — *ISS*_*temporal*_). To test for effects larger than a minimum relevant threshold, (Δ = 0.2), a two-sided Wilcoxon signed-rank threshold test was performed by combining one-sided tests for (*D* > Δ) and (*D* < —Δ). The reported two-sided P value was 2x the smaller one-sided P value, capped at 1, and P values were Bonferroni-adjusted across stages. Confidence intervals were estimated from the Wilcoxon signed-rank test of the paired differences against zero. Computed ISS values can be found in Supplementary Table 1.

#### Spatial identity re-analysis of developmental atlas

To compare spatial identity structure across development, published optic-lobe atlas Seurat objects from Özel et al. were reanalyzed with the same ISS framework^25^. Atlas cluster annotations were mapped to spatial origin^27^, temporal identity^22^, and Notch identity^22^. Cells with missing annotations or multiple spatial origins (i.e., domain-ignoring types) were removed as there is no lineage information to assign true spatial origins to a cell. For each developmental stage, nearest-neighbor graphs were computed from the first 120 PCA dimensions with 20 nearest neighbors, and ISS was calculated for spatial and temporal labels. These atlas scores were compared with scores from the present L3 multiome dataset to evaluate how spatial and temporal identity segregation changed across developmental stages. Computed ISS values can be found in Supplementary Table 1.

### Differential expression/accessibility analysis

To ameliorate pseudoreplication, differential expression and accessibility were tested with pseudobulk edgeR models^72^. For temporal contrasts, counts were aggregated by cell class, temporal identity, spatial origin, and library; spatial origin within library was retained as the replicate-defining unit so that cells from multiple DGRP lines contributed to each pseudobulk when individual DGRP-line counts were too sparse.

Models included the contrast of interest and experiment as a covariate. Library identity defined the pseudobulk samples, including the two separately generated libraries from the final experiment, but was not entered as an additional covariate. Spatial contrasts were tested in neuroepithelial cells, neuroblasts, ganglion mother cells, and neurons. Temporal contrasts were tested in neuroblasts, ganglion mother cells, and neurons. Differential tests were performed as one-vs-rest contrasts within each cell class.

Lowly detected features were removed with edgeR filterByExpr using a minimum count of 10 and a minimum total count of 15 before model fitting. Differential genes and accessible regions were considered significant at FDR < 0.05. Pseudobulk data were normalized with edgeR TMM normalization and modeled with robust dispersion estimation. Differential expression and accessibility were tested using edgeR quasi-likelihood generalized linear models with glmTreat, with a log2 fold-change threshold of 0.5.

Significant features were counted separately for RNA and ATAC, cell class, contrast, and direction of effect based on the sign of the log fold change. Main DEG and DAR count summaries focused on spatial and temporal contrasts. DEGs and DARs were deduplicated by taking the union across one-vs-rest contrasts within each cell class, so that a gene or region significant in multiple contrasts was counted once.

DEG and DAR counts were visualized as signed bar plots. Significant DARs were converted to genomic intervals and visualized across the dm6 genome with karyoploteR density plots, with zoomed views around selected spatial patterning loci. Unfiltered DEGs can be found in Supplementary Table 2; unfiltered DARs can be found in Supplementary Table 3.

### Published chromatin profiles

H3K27me3 CUT&RUN profiles (GSE121028) from Ahmad and Spens^39^ and cell-type-specific Pc DamID profiles (GSE77860) from Marshall and Brand^40^ were visualized at selected regulator loci. Published dm6 coverage files were downloaded and visualized with pyGenomeTracks using the same genomic intervals as the ATAC-seq tracks.

### Code availability

Implementation of analysis of domain-resolved single-cell multiome data can be found in https://github.com/chenyenchung/Chen_et_al_2026_multiome. Cellpose-SAM workflow can be found in https://github.com/chenyenchung/cellpose_nf. Image quantification scripts and results can be found in: https://github.com/chenyenchung/Chen_et_al_2026_microscopy_quant.

### Data availability

All raw and processed data generated in this study are available at the Gene Expression Omnibus with accession number GSE344177. Spatial ISS analysis in mature neurons was done with publicly available single-cell sequencing datasets was retrieved from the Gene Expression Omnibus (GEO) repository with accession number GSE142787. Publicly available larval brain H3K27me3 CUT&RUN dataset was retrieved from the Gene Expression Omnibus (GEO) repository with accession number GSE121028; publicly available larval brain *Pc* DamID was retrieved from the Gene Expression Omnibus (GEO) repository with accession number GSE77860.

## Extended Data Figures

**Extended Data Figure 1.**
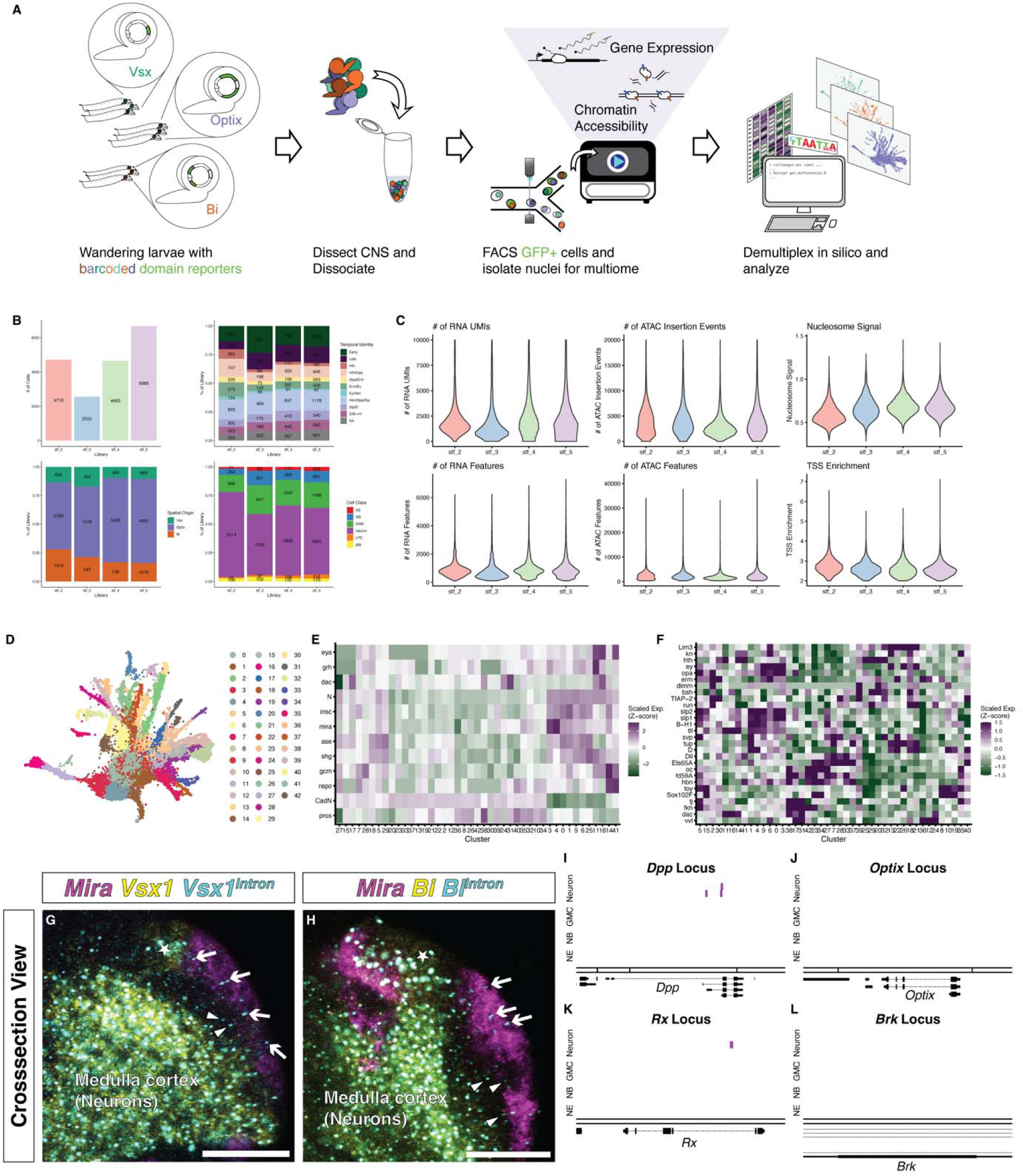
Experimental workflow, quality control, annotation, and additional spatial-patterning loci for single-cell multiome profiling. A, Workflow for spatial-origin multiome profiling. Gal4 reporters expressed in the Vsx, Optix, and Bi neuroepithelial domains drive nuclear GFP, which persists in NB, GMC, and neuronal progenies. GFP-positive cells from reporter lines were pooled across spatial origins within sequencing batches and isolated by FACS. Nuclei were extracted for single-cell RNA/ATAC-seq, and cells were assigned to spatial origin by strain-specific SNP demultiplexing. B, Composition of the profiled dataset across sequencing libraries. Panels show the distribution of recovered cells by library, annotated cell class, temporal identity, and spatial origin. C, Per-cell quality-control metrics for RNA and ATAC modalities across sequencing libraries after filtering. D, Joint RNA–ATAC UMAP embedding colored by unsupervised WNN cluster identity before biological annotation. E, Heatmap of known marker-gene expression across unsupervised WNN clusters used to assign broad cell classes. Marker expression identifies neuroepithelial cells, neuroblasts, ganglion mother cells, neurons, glia, and lamina precursor cells. F, Heatmap of temporal marker expression across WNN clusters used to assign temporal identities. Because individual temporal windows were incompletely resolved in NBs and GMCs, progenitor-stage temporal identities were grouped into early and late cohorts for downstream analyses. G, Cross-sectional view (as in Figure 1B) of *Vsx1* transcript presence relative to *Mira*-expressing NBs (magenta). Cytoplasmic *Vsx1* transcript (yellow) is detected in the neuroepithelium and differentiated neurons, while nascent *Vsx1* transcript (cyan; intron-targeting probes) is detected in the intervening progenitor zone (arrow: transcription puncta in NBs; triangle: transcription puncta in GMCs and newborn neurons; star: neuroepithelial cells). Scale bar: 30μm. H, Cross-sectional view (as in Figure 1B) of *Bi* transcript presence relative to *Mira*-expressing NBs (magenta). Cytoplasmic *Bi* transcript (yellow) is detected in the neuroepithelium and differentiated neurons, while nascent *Bi* transcript (cyan; intron-targeting probes) is detected in the intervening progenitor zone (arrow: transcription puncta in NBs; triangle: transcription puncta in GMCs and newborn neurons; star: neuroepithelial cells). Scale bar: 30μm. I, Locus-level view of spatial DARs at the *Dpp* locus. Each bar represents a unique DAR detected in the indicated cell class; DARs shared across multiple spatial-origin contrasts are collapsed by genomic coordinate. Gene models are shown below the DAR tracks. J, Locus-level view of spatial DARs at the *Optix* locus, plotted as in I. K, Locus-level view of spatial DARs at the *Rx* locus, plotted as in I. L, Locus-level view of spatial DARs at the *Brk* locus, plotted as in I. Unlike the *Vsx1/2* and *Bi* loci shown in Figure 2, these additional spatial-patterning loci in panel I-L show no persistent progenitor-stage domain-specific accessibility comparable to the precise chromatin signature at *Vsx1/2* and *Bi*.

**Extended Data Figure 2.**
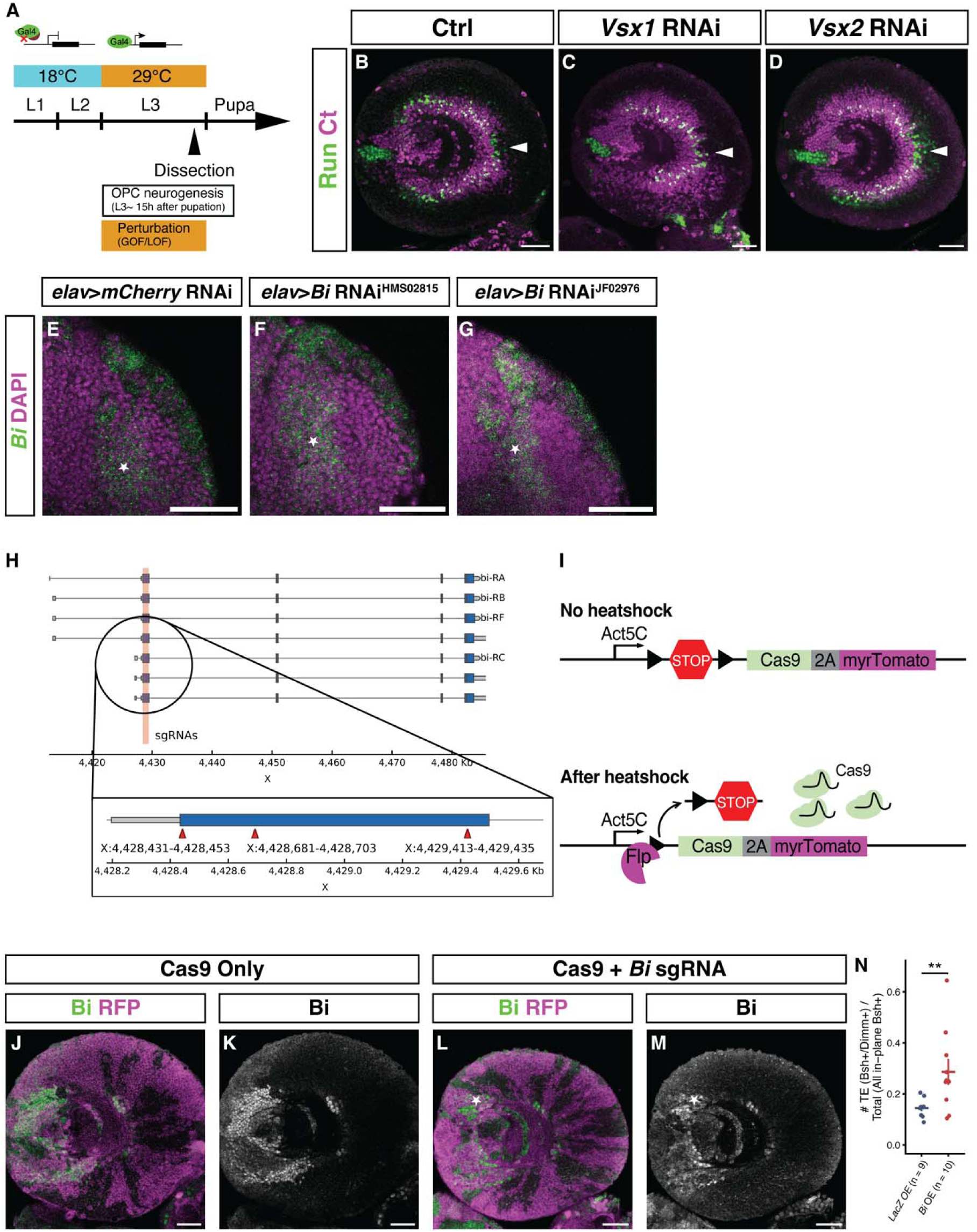
Perturbation strategy and validation for Vsx1/2 and Bi perturbations. A, Schematic of the temperature-sensitive *Gal80* perturbation strategy used for neuronal gain- and loss-of-function experiments. Larvae were maintained at 18°C and shifted to 29°C for the final 48 hours before dissection, corresponding to the third larval instar period when OPC-derived medulla neurons are generated. B-D, Single *Vsx* paralog knockdown in neurons does not eliminate Pm4 neurons. B, Control optic lobe stained for Run and Ct. C, *Vsx1* RNAi. D, *Vsx2* RNAi. White arrowheads indicate persistent Run^+^ Ct^−^ Pm4 neurons after single-paralog knockdown. E–G, Neuronal RNAi does not detectably reduce *Bi* transcript. Cross-sectional views of third-instar larval optic lobes, oriented as in Figure 1B, showing *Bi* transcript detected by HCR-FISH (green) and DAPI (magenta) in an *elav*>*mCherry* RNAi control (E) and animals expressing two independent *UAS-Bi RNAi* transgenes, HMS02815 (F) and JF02976 (G). Stars mark *Bi*-expressing neurons. Neither RNAi line produced a detectable reduction in neuronal Bi transcript. Scale bars: 30μm. H, *Bi* sgRNA design. Whole-locus view of the *Bi* gene with sgRNA target sites indicated; inset shows sgRNAs targeting exon 2, the first coding exon. I, Schematic of the FLP-inducible Cas9 expression cassette used for somatic *Bi* CRISPR. Before heat shock, a transcriptional stop cassette prevents *Cas9-2A-myrTomato* expression from the *Act5C* promoter. Heat-shock-induced FLP excises the stop cassette, enabling Cas9 and myrTomato expression in clones. J-M, Validation of somatic *Bi* CRISPR. J, K, Cas9-only control clone in the posterior optic lobe retaining Bi expression. L, M, Cas9 plus *Bi* sgRNA clones showing broad loss of *Bi* within myrTomato-positive clones. Star indicates a Cas9 plus *Bi* sgRNA clone in which Bi expression persists, likely reflecting incomplete penetrance of somatic CRISPR. Scale bars: 30μm. N, Quantification of TE proportion among Bsh+ cells across the entire optic lobe; n = 9 control and 10 Bi overexpression larvae. TE proportion increased 1.89-fold; quasibinomial regression p = 0.002.

**Extended Data Figure 3.**
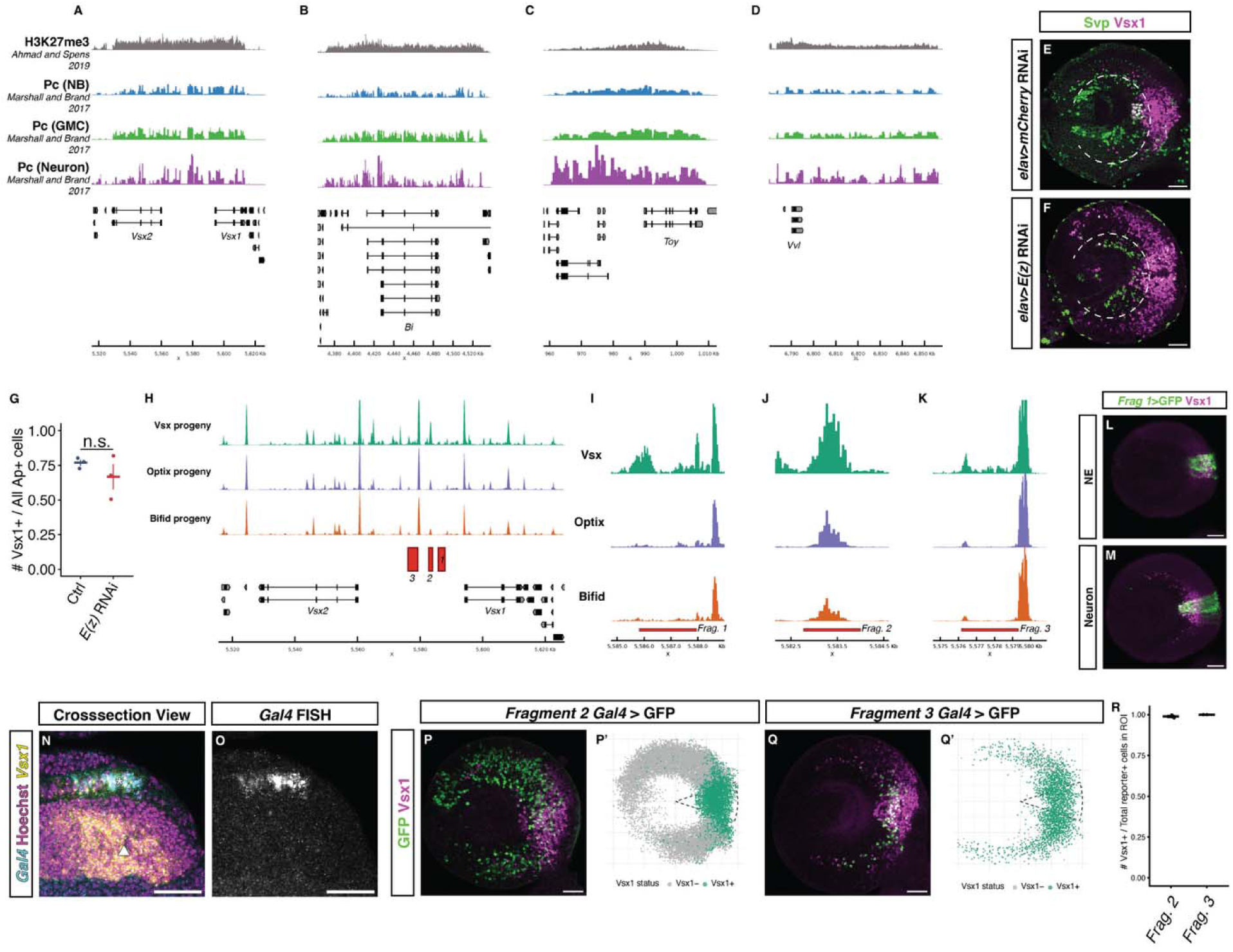
Polycomb occupancy and *Vsx1/2* regulatory-fragment activity support locus-level restriction of neuronal re-expression. A-D, Published H3K27me3 CUT&RUN and cell-type-specific Pc DamID profiles at developmental regulator loci. H3K27me3 CUT&RUN signal from third-instar larval brains (top track, gray) is shown together with Pc DamID coverage in neuroblasts (second track, blue), ganglion mother cells (third track, green) and neurons (bottom track, purple). A, *Vsx1/2* locus. B, *Bi* locus. C, *Toy* locus. D, *Vvl* locus. CUT&RUN data are from Ahmad and Spens, 2019^39^; cell-type-specific DamID data are from Marshall and Brand, 2017^40^. E-F, Vsx1 de-repression under *E(z)* RNAi is biased to later-born neurons. E, Svp (green) and Vsx1 (magenta) in a representative brain section of control (*elav*>*mCherry* RNAi) animals. F, Svp (green) and Vsx1 (magenta) in a representative brain section of animals under neuronal knockdown of *E(z)* (*elav*>*E(z)* RNAi). White dashed lines mark the ring of Svp^+^ neurons born from the early Ey temporal window^22^. G, Fraction of Notch^ON^ neurons (Ap+) that co-express Vsx1 within the conservatively defined Vsx domain (angular coordinate, −π/10 to π/10). Domain-ignoring neurons Mi1, Tm1, Tm2, Tm4, and Tm6 are Notch^ON^ and do not express Vsx1 even when born from the Vsx-domain (also see **Figure 5A-B’**), but Vsx1 is not derepressed in more Vsx-domain Notch^ON^ neurons despite neuronal *E(z)* RNAi. n = 3 control and 3 *E(z)* RNAi larvae. H-K, Neuronal chromatin accessibility at the *Vsx1/2* locus and positions of tested regulatory fragments. H, Neuronal ATAC-seq coverage from the single-cell multiome dataset, separated by Vsx, Optix and Bi spatial origin. The positions of *Fragments 1–3* are indicated. I–K, Enlarged views of *Fragment 1, Fragment 2* and *Fragment 3*, respectively. Coverage was normalized by the library size of each pseudobulk. L-O, *Fragment 1* activity is restricted to the Vsx neuroepithelial domain and is not transcriptionally active in neurons. L, *Fragment 1-Gal4*-driven GFP (green) is present in Vsx1-expressing (magenta) neuroepithelium. M, *Fragment 1-Gal4*-driven GFP is present in Vsx1-expressing anterior medulla neurons. N, Cross-sectional view of *Gal4* transcript detected by FISH (cyan), *Vsx1* transcript (yellow) and Hoechst (magenta). Asterisk: Neuroepithelium; arrowhead: medulla cortex. *Fragment 1-Gal4* transcript is not detected in *Vsx1*-transcribing neurons. O, Grayscale view of the Gal4 FISH channel shown in N. P-Q’, *Fragments 2* and *3* drive broad neuronal reporter expression while retaining high concordance with endogenous Vsx1 within the Vsx domain. P, *Fragment 2-Gal4*-driven GFP and endogenous Vsx1 expression. P’, Segmentation overlay classifying GFP-positive neurons as Vsx1-negative (gray) or Vsx1-positive (green). Q, *Fragment 3-Gal4*-driven GFP and endogenous Vsx1 expression. Q’, Corresponding segmentation overlay. Dashed lines in P’ and Q’ enclose the conservatively defined Vsx domain (angular coordinate, −π/10 to π/10). R, Fraction of GFP-positive neurons that co-express Vsx1 within the conservatively defined Vsx domain for *Fragment 2*- and *Fragment 3-Gal4* reporters. Scale bars in E-F and L-Q: 30μm. n = 3 *Fragment 2* and 3 *Fragment 3* larvae.

**Extended Data Figure 4.**
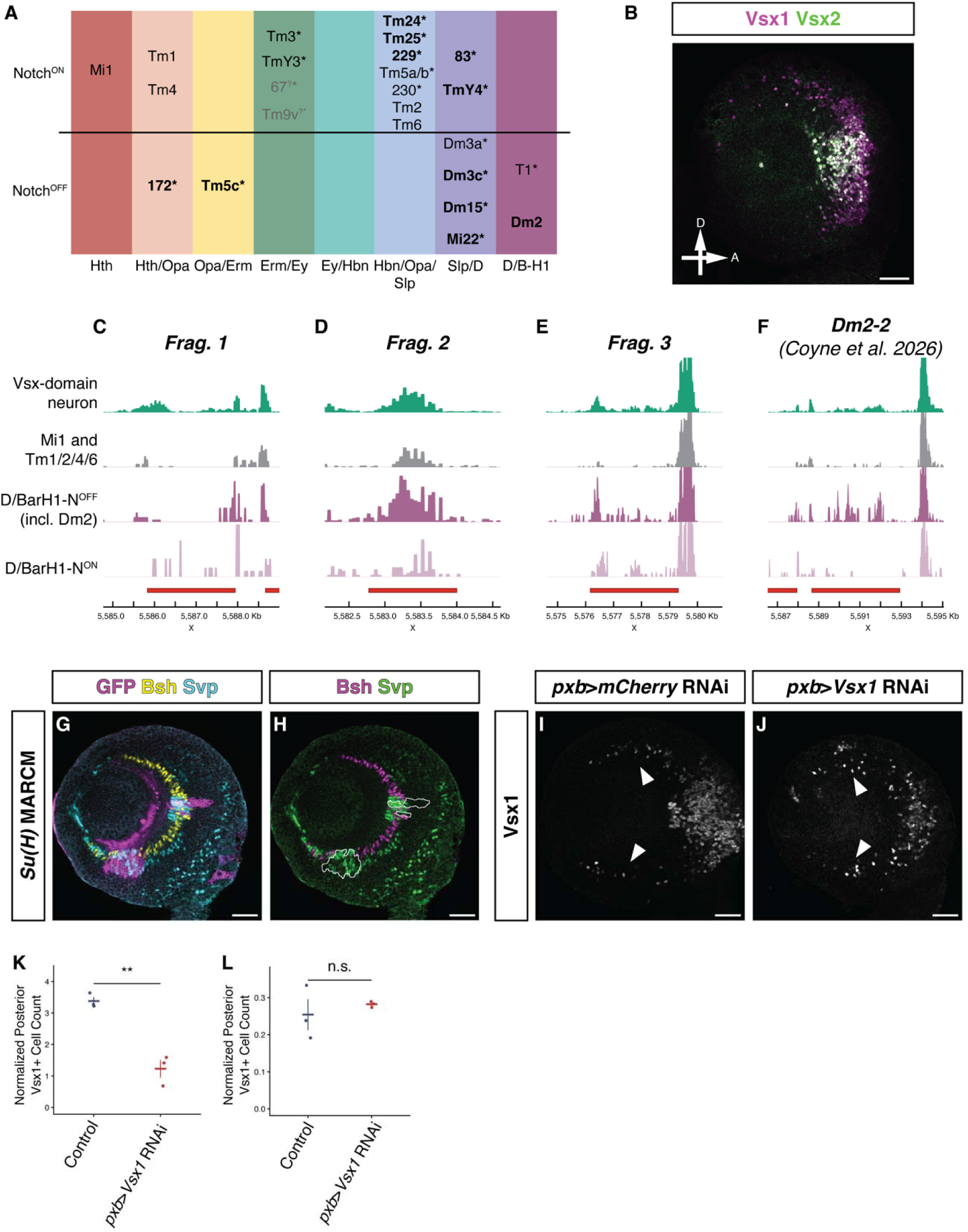
Domain-ignoring type classification, regulatory element accessibility, fate conversion, and domain-independent Vsx1 expression. A, Domain-ignoring types spanning the Vsx-Optix boundary, organized by temporal identity (columns) and Notch status (rows). Bold type indicates types that retain *Vsx1* expression. Asterisks denote types whose *Vsx1* expression status was determined from developmental single-cell RNA-seq^25^. Grayed entries with question marks indicate types that only partially ignore the Vsx-Optix boundary^27^. B, Vsx1 (magenta) and Vsx2 (green) co-expression in Vsx-domain neurons of the third-instar larval optic lobe. Vsx1 and Vsx2 are co-expressed in early- and mid-born neurons, allowing the use of Vsx2 as a proxy for Vsx1 in the *Su(H)* MARCM experiment (Figure 5D-F). Some nascent neurons at the right express Vsx1 but not yet Vsx2, consistent with Vsx2 activation lagging behind Vsx1 in newly born neurons. C-F, Zoomed-in views of pseudobulk ATAC-seq accessibility at the four regulatory regions marked in Figure 5C, using the same neuronal subsets and color scheme. C, *Fragment 1*. D, *Fragment 2*. E, *Fragment 3*. F, *Dm2-2* element^29^. Red bars indicate fragment or element boundaries. G-H, Representative *Su(H)^del47^* MARCM clones showing fate conversion from Mi1 (Bsh+) to Pm3 (Svp+). G, Optic lobe stained for GFP (magenta), Bsh (yellow), and Svp (cyan). H, Clone boundaries outlined in white; Bsh (magenta) and Svp (green). Within clones, Svp-expressing neurons are present but Bsh-expressing Mi1 neurons are absent, consistent with conversion of Mi1 to its Notch^OFF^ sibling fate Pm3. I-J, Vsx-domain neuroepithelial *Vsx1* knockdown (*pxb-Gal4*, which drives specifically in Vsx-domain neuroepithelium) diminishes domain-specific Vsx1 expression while sparing domain-independent Vsx1. I, Control optic lobe (*mCherry* RNAi) showing Vsx1 immunostaining. Arrowheads indicate concentric rings of domain-independent Vsx1-expressing neurons. J, *Vsx1* RNAi. Domain-specific anterior Vsx1 is diminished, while domain-independent Vsx1 expression persists (arrowheads). Anterior is to the right. K, Normalized abundance of Vsx1+ cells in the anterior optic lobe. Vsx1+ neurons in the anterior optic lobe were counted and normalized with the total number of Bsh+ neurons to account for brain size. Welch two-sample t-test p = 0.007. 3 control and 3 Vsx1 NE RNAi larvae. L, Normalized abundance of Vsx1+ cells in the posterior optic lobe. Vsx1+ neurons in the posterior optic lobe were counted and normalized with the total number of Bsh+ neurons to account for brain size. Welch two-sample t-test p = 0.282. 3 control and 3 Vsx1 NE RNAi larvae. Scale bars in B and G-J: 30μm

## Notes

### Competing Interest Statement

The authors have declared no competing interest.

### Summary of Updates

This version has been revised to sharpen and improve presentation. Specifically, the Abstract, Introduction, Results, and Discussion now distinguish persistent locus level chromatin memory from persistence of individual accessible elements. The revised text clarifies that Vsx1 and Bi progenies retain accessibility concentrated at their corresponding spatial factor loci even though the specific accessible regions change across development. We also clarify how PRC2 restricts Vsx1 outside its home domain, and how temporal and Notch identities allow some neuronal types to bypass spatial restriction. New analysis and Extended Data panels show that ectopic Vsx1 expression following neuronal E(z) knockdown is biased toward later born neurons. The Discussion now includes broader comparisons with spatial patterning in the Drosophila ventral nerve cord and vertebrate neural tube and further explains the relationship between spatial bypass and neuronal abundance. Citation was updated, and a Data Availability section was added. The central experimental findings and overall conclusions remain unchanged.

## References

1. Isshiki, T., Takeichi, M. & Nose, A. The role of the *msh* homeobox gene during *Drosophila* neurogenesis: implication for the dorsoventral specification of the neuroectoderm. Development 124, 3099–3109 (1997) 10.1242/dev.124.16.3099.

2. Von Ohlen, T. & Doe, C. Q. Convergence of dorsal, dpp, and egfr signaling pathways subdivides the drosophila neuroectoderm into three dorsal-ventral columns. Dev. Biol. 224, 362–372 (2000) 10.1006/dbio.2000.9789.

3. McDonald, J. A. et al. Dorsoventral patterning in the *Drosophila* central nervous system: the *vnd* homeobox gene specifies ventral column identity. Genes Dev. 12, 3603–3612 (1998) 10.1101/gad.12.22.3603.

4. Doe, C. Q. Molecular markers for identified neuroblasts and ganglion mother cells in the *Drosophila* central nervous system. Development 116, 855–863 (1992) 10.1242/dev.116.4.855.

5. Mellerick, D. M. & Nirenberg, M. Dorsal-ventral patterning genes restrict NK-2 homeobox gene expression to the ventral half of the central nervous system of drosophila embryos. Dev. Biol. 171, 306–316 (1995) 10.1006/dbio.1995.1283.

6. D’Alessio, M. & Frasch, M. msh may play a conserved role in dorsoventral patterning of the neuroectoderm and mesoderm. Mech. Dev. 58, 217–231 (1996) 10.1016/S0925-4773(96)00583-7.

7. Gutjahr, T., Patel, N. H., Li, X., Goodman, C. S. & Noll, M. Analysis of the *gooseberry* locus in *Drosophila* embryos: *gooseberry* determines the cuticular pattern and activates *gooseberry neuro*. Development 118, 21–31 (1993) 10.1242/dev.118.1.21.

8. Skeath, J. B., Zhang, Y., Holmgren, R., Carroll, S. B. & Doe, C. Q. Specification of neuroblast identity in the drosophila embryonic central nervous system by gooseberry-distal. Nature 376, 427–430 (1995) 10.1038/376427a0.

9. Lai, S.-L. & Doe, C. Q. The Fd4 transcription factor translates transient spatial cues in progenitors into long-term lineage identity. eLife 14, RP109188 (2026) 10.7554/eLife.109188.

10. Bhattacharya, A., Rao, H. & Sen, S. Q. Chromatin priming and hunchback recruitment integrate spatial and temporal cues in drosophila neuroblasts. Preprint at 10.7554/eLife.110150.1 (2026).

11. Sen, S. Q., Chanchani, S., Southall, T. D. & Doe, C. Q. Neuroblast-specific open chromatin allows the temporal transcription factor, hunchback, to bind neuroblast-specific loci. eLife 8, e44036 (2019) 10.7554/eLife.44036.

12. Hobert, O. Regulation of terminal differentiation programs in the nervous system. Annu. Rev. Cell Dev. Biol. 27, 681–696 (2011) 10.1146/annurev-cellbio-092910-154226.

13. Hobert, O. & Kratsios, P. Neuronal identity control by terminal selectors in worms, flies, and chordates. Curr. Opin. Neurobiol. 56, 97–105 (2019) 10.1016/j.conb.2018.12.006.

14. Briscoe, J. & Ericson, J. Specification of neuronal fates in the ventral neural tube. Curr. Opin. Neurobiol. 11, 43–49 (2001) 10.1016/S0959-4388(00)00172-0.

15. Malin, J. A., Chen, Y.-C., Simon, F., Keefer, E. & Desplan, C. Spatial patterning controls neuron numbers in the drosophila visual system. Dev. Cell 59, 1132–1145.e6 (2024) 10.1016/j.devcel.2024.03.004.

16. Erclik, T. et al. Integration of temporal and spatial patterning generates neural diversity. Nature 541, 365–370 (2017) 10.1038/nature20794.

17. Erclik, T., Hartenstein, V., Lipshitz, H. D. & McInnes, R. R. Conserved role of the vsx genes supports a monophyletic origin for bilaterian visual systems. Curr. Biol. 18, 1278–1287 (2008) 10.1016/j.cub.2008.07.076.

18. Gold, K. S. & Brand, A. H. Optix defines a neuroepithelial compartment in the optic lobe of the drosophila brain. Neural Dev. 9, 18–34 (2014) 10.1186/1749-8104-9-18.

19. Kaphingst, K. & Kunes, S. Pattern formation in the visual centers of the drosophila brain: wingless acts via decapentaplegic to specify the dorsoventral axis. Cell 78, 437–448 (1994) 10.1016/0092-8674(94)90422-7.

20. Li, X. et al. Temporal patterning of drosophila medulla neuroblasts controls neural fates. Nature 498, 456–462 (2013) 10.1038/nature12319.

21. Bertet, C. et al. Temporal patterning of neuroblasts controls notch-mediated cell survival through regulation of hid or reaper. Cell 158, 1173–1186 (2014) 10.1016/j.cell.2014.07.045.

22. Konstantinides, N. et al. A complete temporal transcription factor series in the fly visual system. Nature 604, 316–322 (2022) 10.1038/s41586-022-04564-w.

23. Suzuki, T., Kaido, M., Takayama, R. & Sato, M. A temporal mechanism that produces neuronal diversity in the drosophila visual center. Dev. Biol. 380, 12–24 (2013) 10.1016/j.ydbio.2013.05.002.

24. Zhu, H., Zhao, S. D., Ray, A., Zhang, Y. & Li, X. A comprehensive temporal patterning gene network in drosophila medulla neuroblasts revealed by single-cell RNA sequencing. Nat. Commun. 13, 1247 (2022) 10.1038/s41467-022-28915-3.

25. Özel, M. N. et al. Neuronal diversity and convergence in a visual system developmental atlas. Nature 589, 88–95 (2021) 10.1038/s41586-020-2879-3.

26. Kurmangaliyev, Y. Z., Yoo, J., Valdes-Aleman, J., Sanfilippo, P. & Zipursky, S. L. Transcriptional programs of circuit assembly in the drosophila visual system. Neuron 108, 1045–1057.e6 (2020) 10.1016/j.neuron.2020.10.006.

27. Simon, F. et al. Spatial, temporal and Notch determination of terminal selector expression controls neuronal cell fate in the Drosophila optic lobe. Nat Neurosci 29, 1068–1078 (2026) 10.1038/s41593-026-02256-6.

28. Suzuki, T., Trush, O., Yasugi, T., Takayama, R. & Sato, M. Wnt signaling specifies anteroposterior progenitor zone identity in the *Drosophila* visual center. J. Neurosci. 36, 6503–6513 (2016) 10.1523/JNEUROSCI.0864-16.2016.

29. Coyne, R. et al. Temporal regulation of a spatial patterning factor in drosophila neurogenesis. Preprint at 10.64898/2026.08.11.744200 (2026).

30. Courgeon, M. & Desplan, C. Coordination between stochastic and deterministic specification in the *Drosophila* visual system. Science 366, eaay6727 (2019) 10.1126/science.aay6727.

31. Kang, H. M. et al. Multiplexed droplet single-cell RNA-sequencing using natural genetic variation. Nat Biotechnol 36, 89–94 (2018) 10.1038/nbt.4042.

32. Mackay, T. F. C. et al. The drosophila melanogaster genetic reference panel. Nature 482, 173–178 (2012) 10.1038/nature10811.

33. Hasegawa, E. et al. Concentric zones, cell migration and neuronal circuits in the *Drosophila* visual center. Development 138, 983–993 (2011) 10.1242/dev.058370.

34. Özel, M. N. et al. Coordinated control of neuronal differentiation and wiring by sustained transcription factors. Science 378, eadd1884 (2022) 10.1126/science.add1884.

35. Holguera, I. et al. Temporal and notch identity determine neuropil targeting depth and synapse location in the fly visual system. Preprint at 10.1101/2025.01.06.631439 (2025).

36. Zhang, Y., Lowe, S., Ding, A. Z. & Li, X. Notch-dependent binary fate choice regulates the Netrin pathway to control axon guidance of Drosophila visual projection neurons. Cell Rep. 42, 112143 (2023) 10.1016/j.celrep.2023.112143.

37. Naidu, V. G. et al. Temporal progression of drosophila medulla neuroblasts generates the transcription factor combination to control T1 neuron morphogenesis. Dev. Biol. 464, 35–44 (2020) 10.1016/j.ydbio.2020.05.005.

38. Islam, I. M., Ng, J., Valentino, P. & Erclik, T. Identification of enhancers that drive the spatially restricted expression of *Vsx1* and *Rx* in the outer proliferation center of the developing *Drosophila* optic lobe. Genome 64, 109–117 (2021) 10.1139/gen-2020-0034.

39. Ahmad, K. & Spens, A. E. Separate polycomb response elements control chromatin state and activation of the vestigial gene. PLOS Genet. 15, e1007877 (2019) 10.1371/journal.pgen.1007877.

40. Marshall, O. J. & Brand, A. H. Chromatin state changes during neural development revealed by in vivo cell-type specific profiling. Nat. Commun. 8, 2271 (2017) 10.1038/s41467-017-02385-4.

41. Chen, Z. et al. A Unique Class of Neural Progenitors in the Drosophila Optic Lobe Generates Both Migrating Neurons and Glia. Cell Reports 15, 774–786 (2016) 10.1016/j.celrep.2016.03.061.

42. Simon, F. et al. Spatial, temporal and notch determination of terminal selector expression controls neuronal cell fate in the drosophila optic lobe. Nat. Neurosci. 29, 1068–1078 (2026) 10.1038/s41593-026-02256-6.

43. Schmitges, F. W. et al. Histone Methylation by PRC2 Is Inhibited by Active Chromatin Marks. Molecular Cell 42, 330–341 (2011) 10.1016/j.molcel.2011.03.025.

44. Yuan, W. et al. H3K36 Methylation Antagonizes PRC2-mediated H3K27 Methylation. J Biol Chem 286, 7983–7989 (2011) 10.1074/jbc.M110.194027.

45. Zhang, Y. et al. Selective bHLH relay factors and modular enhancers decode notch signaling during neuronal diversification in the Drosophila medulla. 2026.06.05.730496 Preprint at 10.64898/2026.06.05.730496 (2026).

46. Treese, M. et al. Regulatory logic of neuronal differentiation in the *Drosophila* visual system. Proc. Natl. Acad. Sci. 123, e2614314123 (2026) 10.1073/pnas.2614314123.

47. Chen, Y.-C. D. et al. Using single-cell RNA sequencing to generate predictive cell-type-specific split-GAL4 reagents throughout development. Proc. Natl. Acad. Sci. 120, e2307451120 (2023) 10.1073/pnas.2307451120.

48. Delás, M. J. et al. Developmental cell fate choice in neural tube progenitors employs two distinct cis-regulatory strategies. Dev. Cell 58, 3–17.e8 (2023) 10.1016/j.devcel.2022.11.016.

49. Isabel Zhang et al. The cis-regulatory logic integrating spatial and temporal patterning in the vertebrate neural tube. bioRxiv 2024.04.17.589864 (2024) 10.1101/2024.04.17.589864.

50. Song, Y., Chung, S. & Kunes, S. Combgap relays wingless signal reception to the determination of cortical cell fate in the drosophila visual system. Mol. Cell 6, 1143–1154 (2000) 10.1016/S1097-2765(00)00112-X.

51. Charest, J. et al. Combinatorial action of temporally segregated transcription factors. Dev. Cell 55, 483–499.e7 (2020) 10.1016/j.devcel.2020.09.002.

52. Cochella, L. & Hobert, O. Embryonic priming of a miRNA locus predetermines postmitotic neuronal left/right asymmetry in C. elegans. Cell 151, 1229–1242 (2012) 10.1016/j.cell.2012.10.049.

53. Matsliah, A. et al. Neuronal parts list and wiring diagram for a visual system. Nature 634, 166–180 (2024) 10.1038/s41586-024-07981-1.

54. Nern, A. et al. Connectome-driven neural inventory of a complete visual system. Nature 641, 1225–1237 (2025) 10.1038/s41586-025-08746-0.

55. Melnattur, K. V. et al. Multiple redundant medulla projection neurons mediate color vision in *Drosophila*. J. Neurogenet. 28, 374–388 (2014) 10.3109/01677063.2014.891590.

56. Morante, J. & Desplan, C. The color-vision circuit in the medulla of drosophila. Curr. Biol. 18, 553–565 (2008) 10.1016/j.cub.2008.02.075.

57. Valentino, P. & Erclik, T. Spalt and disco define the dorsal-ventral neuroepithelial compartments of the developing *Drosophila* medulla. Genetics 222, iyac145 (2022) 10.1093/genetics/iyac145.

58. Boezio, G. L. M. et al. Hierarchical lineage architecture of human and avian spinal cord revealed by single-cell genomic barcoding. Preprint at 10.1101/2025.10.24.684328 (2025).

59. Zhang, I. et al. The cis-regulatory logic integrating spatial and temporal patterning in the vertebrate neural tube. Dev. Cell 60, 3034–3049.e9 (2025) 10.1016/j.devcel.2025.06.029.

60. Dasen, J. S., De Camilli, A., Wang, B., Tucker, P. W. & Jessell, T. M. Hox repertoires for motor neuron diversity and connectivity gated by a single accessory factor, FoxP1. Cell 134, 304–316 (2008) 10.1016/j.cell.2008.06.019.

61. Choi, B. J., Chen, Y.-C. & Desplan, C. Retinal calcium waves coordinate uniform tissue patterning of the *Drosophila* eye. Science 390, eady5541 (2025) 10.1126/science.ady5541.

62. Kuehn, E. et al. Segment number threshold determines juvenile onset of germline cluster expansion in *Platynereis dumerilii*. J. Exp. Zool. Part B 338, 225–240 (2022) 10.1002/jez.b.23100.

63. Pachitariu, M., Rariden, M. & Stringer, C. Cellpose-SAM: superhuman generalization for cellular segmentation. 2025.04.28.651001 Preprint at 10.1101/2025.04.28.651001 (2025).

64. Fleming, S. J. et al. Unsupervised removal of systematic background noise from droplet-based single-cell experiments using CellBender. Nat. Methods 20, 1323–1335 (2023) 10.1038/s41592-023-01943-7.

65. Hao, Y. et al. Integrated analysis of multimodal single-cell data. Cell 184, 3573–3587.e29 (2021) 10.1016/j.cell.2021.04.048.

66. Stuart, T., Srivastava, A., Madad, S., Lareau, C. A. & Satija, R. Single-cell chromatin state analysis with signac. Nat. Methods 18, 1333–1341 (2021) 10.1038/s41592-021-01282-5.

67. Korsunsky, I. et al. Fast, sensitive and accurate integration of single-cell data with harmony. Nat. Methods 16, 1289–1296 (2019) 10.1038/s41592-019-0619-0.

68. Ramírez, F. et al. deepTools2: a next generation web server for deep-sequencing data analysis. Nucleic Acids Res. 44, W160–W165 (2016) 10.1093/nar/gkw257.

69. Lopez-Delisle, L. et al. pyGenomeTracks: reproducible plots for multivariate genomic datasets. Bioinformatics 37, 422–423 (2021) 10.1093/bioinformatics/btaa692.

70. Simpson, E. H. Measurement of diversity. Nature 163, 688–688 (1949) 10.1038/163688a0.

71. Bhargava, T. N. & Uppuluri, V. R. R. Sampling distribution of gini’s index of diversity. Appl. Math. Comput. 3, 1–24 (1977) 10.1016/0096-3003(77)90008-X.

72. Chen, Y., Chen, L., Lun, A. T. L., Baldoni, P. L. & Smyth, G. K. edgeR v4: powerful differential analysis of sequencing data with expanded functionality and improved support for small counts and larger datasets. Nucleic Acids Res. 53, gkaf018 (2025) 10.1093/nar/gkaf018.

